# Antagonistic roles of canonical and alternative RPA in tandem CAG repeat diseases

**DOI:** 10.1101/2022.10.24.513561

**Authors:** Terence Gall-Duncan, Jennifer Luo, Carla-Marie Jurkovic, Laura A. Fischer, Kyota Fujita, David E. Leib, Vanessa Li, Rachel J. Harding, Stephanie Tran, Ran Chen, Hikari Tanaka, Amit L. Deshmukh, Amanda G. Mason, Dominique Lévesque, Mahreen Khan, Stella Lanni, Nozomu Sato, Marie-Christine Caron, Jean-Yves Masson, Gagan B. Panigrahi, Tanya Prasolava, Peixiang Wang, Rachel Lau, Lynette Tippett, Clinton Turner, Albert R. La Spada, Eric I. Campos, Maurice A. Curtis, François-Michel Boisvert, Richard L.M. Faull, Beverly L. Davidson, Hitoshi Okazawa, Marc S. Wold, Christopher E. Pearson

**Author notes:** **Corresponding author:** Christopher E. Pearson. **Contributions** (in order of experiment appearance in manuscript): TGD and VL conducted ddPCR experiments. AGM conducted qRT-PCR. TGD conducted western blots and densitometry quantifications. DEL and BLD provided dissected zQ-175 HD mouse brain. TGD, RL, and PW synthesized repair substrates, TGD and TP generated RPA-deficient functional cell extract, and TGD, JL, and GBP performed *in vitro* repair and binding assays. RJH and CHA perfomed AlphaFold analyses. LAF, RC and MSW purified RPA/Alt-RPA and conducted oligonucleotide binding assays and FRET assays. CMJ, ST, EIC, and F-MB conducted BioID and statistical analysis, DL conducted mass spectrometry analysis, TGD and ST conducted analysis of BioID dataset. TGD, VL, and NS conducted fragment length analysis and SL conducted instability index calculations for SCA1 mouse brains. KF, HT, and HO prepared Rpa1-overexpressing SCA1 mouse brain tissues and conducted mouse immunohistochemistry. LT, CT, MAC, ARLS, and RLMF provided post-mortem patient brain tissues, anonymysed patient information, and sized inherited repeat lengths via PCR. M-CC and J-YM purified FAN1 proteins, ALD performed nuclease assays. TGD, MSW, and CEP conceived of, designed, and coordinated experiments. TGD, JL, KF, LAF, C-MJ, ALD, and GBP prepared figures. TGD, MSW, and CEP wrote the manuscript, J-YM, DEL, BLD, F-MB, EIC, RLMF, MAC, ARL, and HO provided edits and comments. All authors approved of the final text, data and figures.

## Abstract

Tandem CAG repeat expansion mutations cause >15 neurodegenerative diseases, where ongoing expansions in patients’ brains are thought to drive disease onset and progression. Repeat length mutations will involve single-stranded DNAs prone to form mutagenic DNA structures. However, the involvement of single-stranded DNA binding proteins (SSBs) in the prevention or formation of repeat instability is poorly understood. Here, we assessed the role of two SSBs, canonical RPA (RPA1-RPA2-RPA3) and the related Alternative-RPA (Alt-RPA, RPA1-RPA4-RPA3), where the primate-specific RPA4 replaces RPA2. RPA is essential for all forms of DNA metabolism, while Alt-RPA has undefined functions. RPA and Alt-RPA are upregulated 2- and 10-fold, respectively, in brains of Huntington disease (HD) and spinocerebellar ataxia type 1 (SCA1) patients. Correct repair of slipped-CAG DNA structures, intermediates of expansion mutations, is enhanced by RPA, but blocked by Alt-RPA. Slipped-DNAs are bound and melted more efficiently by RPA than by Alt-RPA. Removal of excess slipped-DNAs by FAN1 nuclease is enhanced by RPA, but blocked by Alt-RPA. Protein-protein interactomes (BioID) reveal unique and shared partners of RPA and Alt-RPA, including proteins involved in CAG instability and known modifiers of HD and SCA1 disease. RPA overexpression inhibits rampant CAG expansions in SCA1 mouse brains, coinciding with improved neuron morphology and rescued motor phenotypes. Thus, SSBs are involved in repeat length mutations, where Alt-RPA antagonistically blocks RPA from suppressing CAG expansions and hence pathogenesis. The processing of repeat length mutations is one example by which an Alt-RPA↔RPA antagonistic interaction can affect outcomes, illuminating questions as to which of the many processes mediated by canonical RPA may also be modulated by Alt-RPA.

## INTRODUCTION

Over 69 neurodegenerative, neurological, and neuromuscular diseases are caused by expansions of gene-specific tandem repeat DNA sequences^1^. At least 15 are caused by (CAG)•(CTG) repeat expansions, including Huntington disease (HD) and many spinocerebellar ataxias (SCAs) including types 1 and 3 (SCA1 and SCA3). In HD and SCA1, disease manifests when the repeat expands to ≥36 or ≥39, respectively, while unaffected lengths are shorter and genetically stable. Inherited expansions continue to somatically expand in affected tissues as patients age, supporting the concept that ongoing expansions drive disease onset and progression^2–4^. Since disease age-of-onset (AOO), progression, and severity are all associated with repeat size, somatic repeat expansion could be a major driver of pathogenesis with age.

Human genome-wide association (GWA) data for HD, SCA1, and other repeat expansion diseases have validated three related modifiers of AOO, strengthening a connection of somatic instability to disease^5–10^. The first two modifiers, repeat tract length and purity, enhance and slow CAG expansions, respectively, by enhancing or diminishing the formation of slipped-CAG DNAs which are intermediates of expansion mutations. The third modifier, DNA repair protein variants, were identified as modifiers of AOO and disease progression in the repeat expansion HD, SCA1-3, SCA6, SCA7, SCA17, myotonic dystrophy type 1 (DM1), and X-linked dystonia-parkinsonism (XDP)^10–15^. Importantly, each of these disease-modifying DNA repair proteins (FAN1, MSH3, MLH1, PMS2, PMS1, LIG1, and POLD1)^16^, are also modifiers of somatic CAG expansions in brains of HD mice, likely through their processing of slipped-CAG structures. These strong human and mouse data support a causal link between repeat instability with disease onset and progression. This link is further strengthened by the identification of naturally-occurring variants of the human and murine MSH3/Msh3 that favor or disfavor somatic CAG expansions in HD and DM1 patients and mice^17^, which parallels with their modfying disease onset and progression^11^. While the GWA studies have validated the involvement in humans many of the proteins known to be involved in somatic CAG expansions, they have not identified all such factors. For example, MSH2 and MLH3 are equally crucial to somatic CAG expansions as their mismatch repair protein binding partners MSH3 and MLH1, but only the latter two were identified in the GWA screens^11,18–20^. Thus, it is possible that other DNA repair proteins not identified by GWAS may be involved in the formation of CAG expansions. Due to the formation of unusual DNA structures from single-stranded tandem repeats^21,22^, clear candidates are the single-strand DNA binding (SSB) proteins, essential to all pathways of DNA metabolism.

Slipped-CAG DNA structures formed at expanded repeats are mutagenic intermediates of somatic expansions and arise by misaligned base-pairing (strand-slippage). Strand-slippage can occur when DNA is unwound into single-strands during transcription, replication, repair, or recombination^8,21,23–25^. For example, transcription across expanded repeats in post-mitotic neurons enhances somatic expansions^26–29^. Slipped-DNAs have also been identified at the expanded (CAG)•(CTG) repeat of the mutant *DMPK* gene in DM1 patient tissues, with tissue-specific levels correlating directly with increased somatic repeat expansions^30^. Recent evidence supports slipped-DNAs as critical intermediates of instability. Earlier, we showed that small-molecule targeting of mutagenic slipped-CAG structures inhibits expansions and elicits contractions of the mutant CAG tract *in vivo* in HD mouse brains^29^. Thus, understanding how slipped-DNAs are processed to expansions is important for therapeutic development.

One of the most important and well characterized SSBs is canonical RPA, a heterotrimeric complex composed of RPA1 (70 kDa), RPA2 (32 kDa), and RPA3 (15 kDa) (Figure 1A). RPA is essential for life, ubiquitously expressed, and mediates virtually every process involving ssDNA (replication, repair, recombination, transcription, etc.)^31–34^. This includes DNA unwinding, melting of unusual DNA structures, reannealing of DNAs, binding and protecting single-strand DNA, and protein recruitment to DNA^35–39^. RPA is highly conserved, with all eukaryotes containing homologs of each subunit^34^. Considering the likely involvement, it is surprising that only two studies focused on the impact of single-strand binding proteins on repeat instability^40,41^.

**Figure 1:**
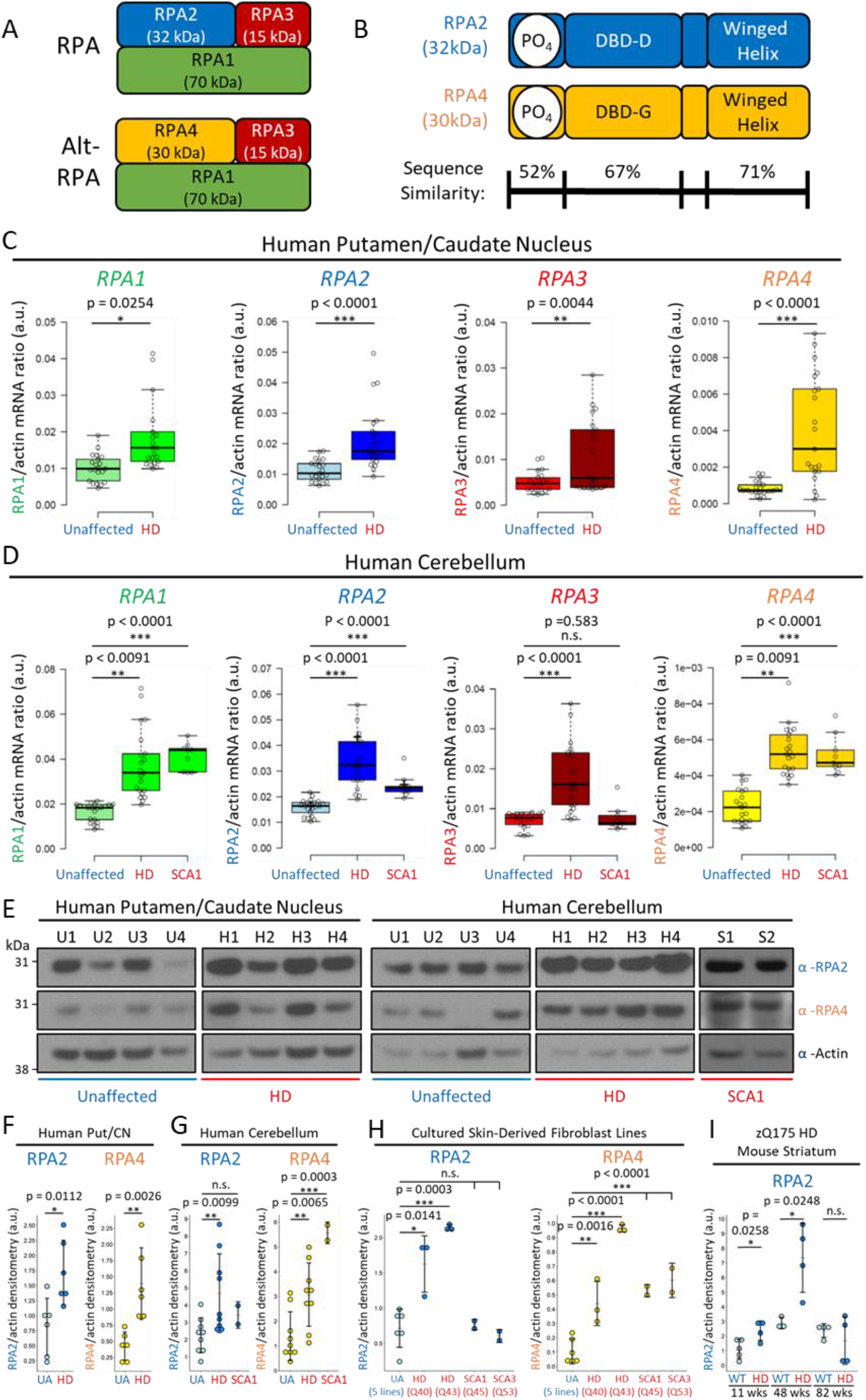
RPA1, RPA, RPA3, and RPA4 are upregulated at the RNA and protein level in patient brain tissues, cell lines, and mouse brain tissues. A) canonical RPA complex composed of subunits RPA1, RPA2, and RPA3, vs alternative RPA complex composed of RPA1, RPA4, and RPA3. B) sequence differences between RPA2 and RPA4 reveal similar DNA binding domains (DBD; protein-DNA interactions) and winged helix domains (protein-protein interactions), relative to a less similar N-terminal site amenable to phosphorylation. C) ddPCR data of all RPA subunits from the striatum of HD patients. n = 7 individuals per group per tissue, 3 replicates per subject. Statistics: unpaired student t-test comparing means. D) ddPCR data of all RPA subunits from the cerebellum of HD and SCA1 patients. n = 7 HD and unaffected individuals per group per tissue, and 3 SCA1 patients per group per tissue, 3 replicates per person. E) Representative western blots for HD and SCA1 patient striatum and cerebellum relative to unaffected control tissues, probing for RPA2 and RPA4 expression levels. Actin loading control. F-I) densitometric quantification of RPA2 and RPA4 levels as a ratio of actin loading in the striatum, cerebellum, cultured fibroblasts, and zQ175 HD mouse striatum. Statistics: unpaired student t-test comparing means.

In addition to canonical RPA, primates also express an alternative form of RPA, known as Alt-RPA. Alt-RPA is composed of RPA1-RPA4-RPA3 (Figure 1A). Thus, Alt-RPA differs from RPA via the swap of RPA2 (32 kDa) to RPA4 (30 kDa). Either RPA2 or RPA4, but not both simultaneously, interacts with RPA1 and RPA3 to form canonical RPA or Alt-RPA heterotrimeric complex, respectively^42,43^. *RPA4*, at Xq21.33, a paralog of *RPA2*, is present only in primates and a limited number of placental mammals (excluding mice)^42–44^. RPA4 has 47% and 63% amino acid sequence identity and similarity to RPA2, with similar domain organization (Figure 1B)^43^. In contrast to canonical RPA, functionally characterized by >3000 studies, surprisingly little is known about Alt-RPA, being limited to just 6 studies^42–47^. RPA4 is less abundant than RPA1 and RPA2, is present in non-proliferating tissues, with low expression in brain^42,48^. RPA4 is reduced in cancers, and cannot support cell cycle progression, consistent with a role in genome maintenance in non-proliferating cells^42,44,47^.

Here, we assessed RPA and Alt-RPA levels in human brains and their roles in somatic CAG repeat instability. We observe RPA and Alt-RPA are upregulated 2- and 10-fold, respectively, in HD and SCA1 patient brains relative to age- and sex-matched non-affected controls. Further, our data show that RPA enhances *in vitro* repair of slipped-CAG repeats, while high levels of Alt-RPA can block slipped-DNA repair. Both RPA and Alt-RPA can bind slipped-DNAs, but only RPA can efficiently melt them. FAN1 nuclease (a modifier of HD age-of-onset) cleavage of slipped-CAG structures is enhanced by RPA, but inhibitied by Alt-RPA. BioID of each RPA subunit revealed unique and shared association with proteins important for somatic repeat instability. Over-expressing the murine Rpa1 subunit ablates spontaneous somatic CAG expansions *in vivo* in brains of SCA1 mice. Additionally, we find that RPA-mediated inhibition of somatic expansions coincides with reduced levels of disease biomarkers including the DNA damage response markers γ-H2AX and 53BP1, diminished ubiquitin-positive mutant polyQ Ataxin-1 aggregation in striatal neurons and improved motor functions. In sum we provide evidence that the SSB proteins RPA/Alt-RPA, and their relative ratios, modulate CAG repeat instability.

## RESULTS

### RPA and Alt-RPA are upregulated in HD and SCA1 patient brains

We first quantified RNA and protein levels of *RPA1*, *RPA2*, *RPA3*, and *RPA4* in post-mortem brains from HD and SCA1 patients along with age- and sex-matched unaffected individuals. We assessed brain regions that show varying disease vulnerability and somatic CAG expansions: in HD patients, the striatum (dorsally comprised of the putamen and caudate nucleus) is highly degenerated, the cerebellum is moderately degenerated, and the frontal pole does not degenerate^49–51^. In SCA1, the cerebellum is the most highly degenerated brain tissue, with evidence of degeneration in the striatum in late stage disease^52–54^. In HD and SCA1 the levels of somatic CAG expansions are very high in the striatum and negligible in the cerebellum^2–4^. Brain regions per cohort, patient information, and disease staging, are detailed in Table 1.

| Huntington Disease patients: |  |  |  |  |  |  |  |  |
| --- | --- | --- | --- | --- | --- | --- | --- | --- |
| Case # | In text code | HD grade | Age at disease onset | Age at death | M/ F | Post mortem delay (hr) | Cause of death | HTT CAG repeat lengths |
| HC77 | H7 | HD-4 | 29 | 53 | F | 9 | Pneumonia | 17/54 |
| HC133 | H6 | HD-2 | 48 | 65 | M | 14 | Renal failure | 17/43 |
| HC147 | H5 | HD-3 | 30 | 64 | M | 18 | Pulmonary thromboembolus | 27/42 |
| HC148 | H4 | HD-3 | 39 | 63 | M | 16 | Pulmonary embolism | 22/43 |
| HC150 | H3 | HD-2 | 51 | 64 | F | 21 | Bronchopneumonia | 22/42 |
| HC161 | H2 | HD-2 | n/a | 70 | F | 8 | HD | 17/42 |
| HC132 | H1 | HD-1 | 31 | 32 | M | 14 | Submandibular squamous cell carcinoma | 17/47 |
| Unaffected individuals: |  |  |  |  |  |  |  |  |
| H170 | U7 | control | - | 60 | M | 17 | Ischaemic heart disease | 10/17 |
| H174 | U6 | control | - | 59 | M | 24.5 | Aortic aneurysm | 17/18 |
| H181 | U5 | control | - | 78 | F | 20 | Aortic aneurysm | 18/19 |
| H194 | U4 | control | - | 68 | M | 22.5 | Coronary atherosclerosis | 17/19 |
| H209 | U3 | control | - | 48 | M | 23 | Ischaemic heart disease | 17/20 |
| H239 | U2 | control | - | 64 | M | 15.5 | Ischemic heart disease | 13/15 |
| H177 | U1 | control |  | 22 | M | 21 | Asphyxia | 11/23 |

Within the striatum, transcripts for all four RPA genes were significantly upregulated in HD patients relative to unaffected individuals (Figure 1C) as assessed via droplet digital PCR (ddPCR). *RPA1* was increased ~1.5-fold (p=0.0254), *RPA2* was increased ~2-fold (p<0.0001), *RPA3* was increased ~1.2-fold (p=0.0044), and *RPA4* exhibited striking upregulation of ~5-6-fold (p<0.0001) (Figure 1C). These findings were validated by real-time quantitative reverse transcription PCR (qRT-PCR) in a separate cohort of HD patients relative to unaffected individuals, and similar upregulations were observed for *RPA2* (~1.5-fold, p=6.6e^−6^) and *RPA4* (~8-fold, p=9.5e^−5^) (Supplementary Figure S1A). Upregulation at the protein level was also observed at similar fold increases in the striatum, for RPA2 (~2-fold, p=0.0112) and RPA4 (~3-4-fold, p=0.0026) (Figure 1E, Supplementary Figure S1B; quantified in Figure 1F). These findings were also seen in HD patient fibroblast cell lines (Q43 and Q40) with ~2-3-fold upregulation of RPA2 and ~5-10-fold upregulation of RPA4 observed relative to 5 control lines (Supplementary Figure S1I; quantified in Figure 1H).

The cerebellum also showed RPA subunit RNA upregulation via ddPCR, with differences observed between HD and SCA1 patients relative to unaffected individuals (Figure 1D). *RPA1* was upregulated in HD (~1.5-fold, p<0.0091) and SCA1 (~2-fold, p<0.0001). *RPA3* was upregulated ~2-fold in HD (p<0.0001), but not in SCA1 (p=0.583) (Figure 1D). Similarly, while *RPA2* was upregulated in HD (~2-fold, p<0.0001) and SCA1 (p<0.0001), the magnitude of upregulation was mild in the SCA1 cerebellum (~1.2-fold) (Figure 1D). *RPA4* was moderately upregulated by ~2.5-fold in both HD (p=0.0091) and SCA1 (p<0.0001) (Figure 1D). qRT-PCR of the separate cohort confirmed similar upregulation of *RPA2* (~2-fold, p=8.6e^−7^) and *RPA4* (~6-fold, p=2.8e^−7^) (Supplementary Figure S1A).

Protein upregulation was also evident in HD patient cerebellum for both RPA2 (~2.5-fold, p=0.0099) and RPA4 (~2.5-fold, p=0.0065) (Figure 1E, Supplementary Figure S1B; quantified in Figure 1G). In contrast, in SCA1 patient cerebellum, while ~4.5-fold upregulation of RPA4 protein was observed (p=0.0003), RPA2 protein was not upregulated (p=0.1624) despite *RPA2* upregulation at the RNA level (Figure 1E; quantified in Figure 1G). Consistent with these findings, RPA2 did not exhibit upregulation in SCA1 (p=0.8524) and SCA3 (p=0.4642) patient fibroblasts, despite upregulations of RPA4 (~5-6-fold, p<0.0001) in both cell lines (Supplementary Figure S1I; quantified in Figure 1H).

In contrast to the other brain regions, the frontal pole of HD patients showed insignificant upregulation of *RPA1* (p=0.2333), and mild upregulation with limited spread between individuals for *RPA2* (~1.5-fold, p=0.0006), *RPA3* (~1.2-fold, p=0.0015), and *RPA4* (~2-fold, p=0.039) (Supplementary Figure S1C), via ddPCR. However, neither RPA2 nor RPA4 were significantly upregulated in the frontal pole at the protein level (Supplementary Figure S1D; quantified in Supplementary Figure S1E).

Cumulatively, we conclude that Alt-RPA is highly upregulated in HD and SCA1 patient brains, while canonical RPA is mildly upregulated. Moreover, brain region expression patterns corelated with degenerative vulnerability in HD: RPA4 showed striking upregulation and RPA2 showed mild upregulation in the striatum (the most degenerated brain region), mild upregulations of both RPA2 and RPA4 were observed in the cerebellum (a less degenerated tissue), and no change in RPA2 and RPA4 levels in the frontal pole (which does not undergo neurodegeneration). While brain region specificity could not be assessed in the SCA1 patients due to limited samples, RPA4 was highly upregulated while RPA2 was not upregulated within the cerebellum (the most degenerated brain region in SCAs). These data support that upregulation of Alt-RPA expression (or increased ratios of Alt-RPA to RPA) may coincide with disease-specific repeat expansions and/or neurodegeneration^2–4^.

### RPA and Alt-RPA expression in HD tracks with disease stage

We next assessed if *RPA2* and *RPA4* transcript levels fluctuated with i) the age at which patients succumbed to their disease, ii) the inherited CAG length, and/or iii) the neuropathological grading (Vonsattel)^55^ in HD patients. RPA2 showed a slight negative correlation with age (R^2^=0.53) in the striatum of HD patients. No trend was observed for RPA2 expression in the cerebellum and RPA4 expression in both the striatum and the cerebellum (Supplementary Figure S1J). Furthermore, no correlation was observed for RPA2 and RPA4 expression with inherited repeat length within any tissues (Supplementary Figure S1K). We therefore conclude that RPA2 and RPA4 expression level fluctuations are not linked with age or inherited CAG length.

Interestingly, negative correlations were observed for RPA2 and RPA4 expression with neuropathological grading within the striatum of HD patients - with RPA2 and RPA4 levels decreasing as grades increased (Supplementary Figure S1L). These findings may be reflective of the level of neurodegeneration observed within these tissues, such that upregulation of RPA2 and RPA4 is not observed as more transcriptionally dysregulated neurons progressively die at higher grades. In contrast, RPA2 expression levels increased with grade in the HD cerebellum, while RPA4 levels did not change. Since the cerebellum is less degenerated in HD, this supports a model whereby increased ratios of RPA to Alt-RPA reflect lower levels of neurodegeneration (Supplementary Figure S1L).

We next assessed Rpa2 protein levels in striatum and cerebellum of the zQ175 HD mouse model prior to disease onset (11 weeks), in mid-disease (48 weeks), and in late disease (82 weeks)^56^. Since mice contain a non-functional *Rpa4* pseudogene, Rpa4 expression was not assessed. Rpa2 expression in the zQ175 HD mouse striatum was significantly upregulated compared to wildtype controls at 11 weeks (~2-fold, p=0.0258) and 48 weeks (~7-fold, ap=0.0248) before dropping to wild-type levels at 82 weeks (p=0.4429) (Supplementary Figure S1F; quantified in Figure 1I). The reduction in RPA2 expression observed between 48-week-old and 82-week-old zQ175 mice is similar to trends observed in patients with age (Supplementary Figure S2A). Unlike HD patients, Rpa2 protein levels in the zQ175 HD mouse cerebellum did not differ relative to wild-type at any age group (Supplementary Figure S1G; quantified in Supplementary Figure S1H). Unlike HD patients, the zQ175 HD mouse model exhibits little to no neurodegeneration with age, suggesting that the drop in *Rpa2* expression at 82 weeks is unlikely to be the result of disease-associated neurodegeneration.

Temporally, CAG repeat expansions tend to increase with age in human HD and mouse HD striatal brain regions^2,4,17,57,58^. Interestingly, the onset of striatal CAG expansion parallels the timing of RPA2 upregulation in HD mice striata, suggesting that shifting levels of RPA could be linked with fluctuating rates of somatic expansions. To consider the functional roles of RPA and Alt-RPA in mediating somatic repeat instability, and how their dysregulated expression levels may affect this, we next assessed for structural differences between the RPA and Alt-RPA which could indicate differential functions.

### Subtle structural differences could underly differential RPA and Alt-RPA functions

Considering the high degree of sequence similarity between RPA2 and RPA4 (Figure 1B), we hypothesize that potential differential functions of RPA and Alt-RPA are likely due to differences in 3D structure. In the absence of any experimentally determined structures of RPA4 in isolation or in complex with other proteins, the AlphaFold predicted RPA4 model structure was analysed (Supplementary Figure S2A-C)^59–61^. RPA4 has 67% sequence identity with RPA2 (Supplementary Figure S2D), is predicted to also have a DNA binding domain (DBD) and a winged-helix domain, like RPA2. The Alt-RPA hetero-trimerization core was modelled by superposition of RPA4 DBD-G onto RPA2 DBD-D (Supplementary Figure S3E), revealing subtle changes in the surface electrostatics compared to the canonical RPA complex which might be expected to elicit differential abilities to interact with nucleic acids (Supplementary Figure S2F). Furthermore, other suble differences may also contribute to differential subunit-subunit and subunit-protein interactions to affect functional outcomes. For example, although most key RPA1 and RPA3 interface residues in RPA2 (inter-protein distance <3.5A) are conserved in RPA4, the RPA1-interacting FKIM (Phe-Lys-Ile-Met) motif in RPA2 (Supplementary Figure S2G) is not conserved in RPA4. We hypothesize that subtle changes in the inter-protein interfaces, possible small changes in the complex electrostatics, and/or other structural differences of RPA2 and RPA4 could promote differences in RPA and Alt-RPA activity. Based on these findings, we next probed for how these structural differences could impact *in vitro* repair of slipped-CAG/CTG structures, intermediates of somatic CAG expansions.

### Characterizing repair of nick-in-repeat slipped-DNA repair substrates

RPA is required for processing partially single-stranded DNA intermediates. *In vitro* repair of slipped-DNAs by human cell extracts has previously been used to elucidate the roles of DNA repair proteins in processing slipped-DNAs^62–66^. We assessed the roles of RPA and Alt-RPA in *in vitro* slipped-DNA repair. We used CAG/CTG slipped-DNA intermediates with nicks located in the repeat (nick-in-repeat) or flanking sequences (nick-in-flank). Nicks in the repeat tract may arise during repair of DNA damage, gap-filling, or polymerase slippage. Nicks located within the repeat at the extreme end of the repeat tract can generate free-end repeat strands which experience greater degrees-of-freedom, thereby allowing them to interconvert between heterogenous random coil slipped conformations (Figure 2A) Nicks in the repeat might incur increased opportunities for end-fraying slippage events, where the free-end of the repeat translationally realigns (Figure 2A). In contrast, when the nick is in the flanking regions, the excess repeats are anchored by ~60 basepairs of duplex non-repeat flanks and assume a defined hairpin slip-out with a single 3-way junction (Figure 2A). In this way, nick-in-repeat Δ(CAG)20 slip-outs [(CAG)50•(CTG)30] are molecularly identical to nick-in-flank slipped-DNAs, differing only in the nick location and secondary structure (Supplementary Figure S3A and 3B). As expected, nick-in-repeat slipped-DNAs are conformationally distinct from nick-in-flank slipped-DNA, as evidenced by its increased electrophoretic migration relative to nick-in-flank substrates (Figure 2, Supplementary Figure S3C, compare lanes 1 between panel A and B, see black versus white arrowheads, see detailed Supplementary Text). Previously, heterogenous random-coil and hairpin conformations of the same sequences, were demonstrated to have differential electrophoretic mobilities^67–73^.

**Figure 2:**
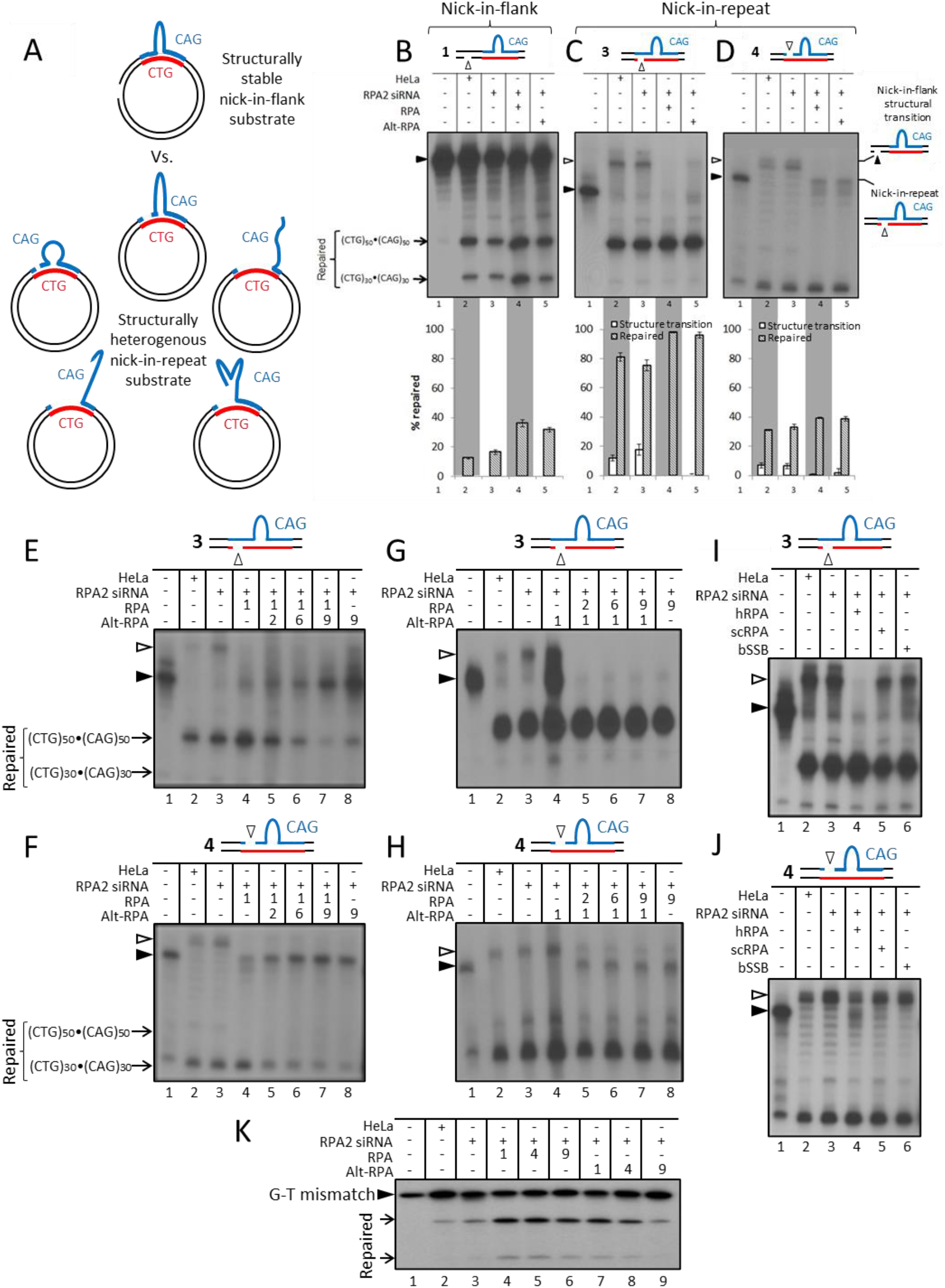
RPA and Alt-RPA both enhance repair at low concentrations, but Alt-RPA inhibits repair at higher concentrations. A) Nick-in-repeat substrates have a nick within the first repeat of the CTG/CAG repeat tract, allowing the repeat tract on the nicked strand to slip and slide along the continuous strand. Thus nick-in-repeat substrates have high structural variability and increased single-stranded DNA formation in both strands. B-D) Molecularly identical slipped-DNAs with varying nick locations are indicated by schematics. Repair reactions were performed with cold dNTPs, repeat-containing fragments released by EcoRI/HindIII digestion, and repair products visualized by Southern blotting following resolution on native 4% PAGE. Reaction products for panels A, B, and C were run on the same gel. As indicated substrates were processed by HeLa or HeLa cells knocked down for RPA (RPA2 siRNA treated), and the last two lanes of each panel have added recombinant RPA or Alt-RPA. Starting SI-DNA bands and repair products are indicated with a black triangle, a slower migrating species is indicated by white triangle/hollow arrow heads. Lanes 1 of each panel shows starting SI-DNA substrates without processing. Subsequent lanes show each substrate processed by 25 μg of HeLa cell extract (lane 2), 25 μg RPA-deficient extract HeLa cell extract alone (lane 3), or supplemented with 600 ng RPA (lane 4), or 600 ng Alt-RPA (lane 5). The starting SI-DNA substrates are indicated by black triangles. The slow-migrating species that migrate above the starting substrate after processing by cell extracts are indicated by white triangles/hollow arrow heads. Repaired products of (CTG)50•(CAG)50 and (CTG)30•(CAG)30 are indicated. Repair efficiencies were calculated for three to six replicates and adjusted for starting background. Graph shows repaired material (cross-hatched bars) and processed material that migrates above the starting SI-DNA (white bars). E-H) High levels of RPA enhance, while high levels of Alt-RPA block, slipped-CAG repair. Repair reactions were performed with cold dNTPs, the repeat-containing fragments released by EcoRI/HindIII digestion, and repair products visualized by South Blotting following resolution on native 4% PAGE. A plus sign indicates the addition of the indicated component. Starting unprocessed SI-DNAs and repair products are indicated. Lane 1 shows starting unprocessed SI-DNA substrate. Lane 2 shows substrate processed by 25 μg HeLa cell extract. Lane 3 shows substrate processed by 25 μg RPA-deficient HeLa cell extract (RPA-/-). Lane 4 shows substrate processed by RPA-deficient extract supplemented by 600 ng RPA or Alt-RPA. Lane 5-7 shows processing by RPA-deficient extract supplemented by 600 ng RPA or Alt-RPA and 1.2 μg (1:2), 3.6 μg (1:6), or 6 μg (1:10) RPA or Alt-RPA, respectively. Lane 8 shows substrate processed by RPA-deficient extract supplemented by 6 μg RPA or Alt-RPA. I-J) RPAs slipped-DNA repair functionality cannot be replaced by bacterial SSB or yeast RPA. Repair reactions were performed with cold dNTPs, the repeat-containing fragments released by EcoRI/HindIII digestion, and repair products visualized by South Blotting following resolution on native 4% PAGE. Products in both panels were run on the same gel. A plus sign in the table indicates the addition of the indicated component. Starting SI-DNA bands and repair products are indicated with a black triangle, a slower migrating species is indicated by a hollow arrow head. Lane 1 shows starting unprocessed SI-DNA substrate. Lane 2 shows substrate processed by 25 μg of HeLa cell extract. Lane 3 shows substrate processed by 25 μg of RPA-deficient HeLa cell extract (RPA-/-). Lanes 4 to 6 show RPA-deficient cell extract supplemented with 600 ng of the indicated protein. Repaired products of (CTG)50•(CAG)50 and (CTG)30•(CAG)30 are indicated.

*In vitro* repair of the nick-in-repeat slip-outs was compared to nick-in-flank slip-outs (Supplementary Figure S3A). *In vitro* repair of nick-in-repeat and nick-in-flank Δ(CAG)20 slip-outs [(CAG)50•(CTG)30] by HeLa cell extract yielded the correct DNA repair products expected from nick-directed slip-out repair (Figure 2B-D, Supplementary Figure S3C, compare lanes 1 and 2 in each panel). In both cases the nicked (CTG)30 strand was repaired and the continuous (CAG)50 strand was used as a template for repair, yielding the duplex product of (CTG)50•(CAG)50 (Figure 2B-D, Supplementary Figure S3C, compare lanes 1 and 2 in each panel). Qualitative differences in the reaction products were evident between nick-in-flank and nick-in-repeat slip-outs. Repair of nick-in-repeat slipped-DNAs [Δ(CAG)20 slip-outs (CAG)50•(↓CTG)30 or (↓CAG)50•(CAG)30; where nick location is defined by the ↓] yielded a series of products that electrophoretically migrated slower than their starting substrate - and which co-migrated with the starting unrepaired nick-in-flank substrates (Supplementary Figure S3C, compare white arrowheads in lanes 4 and 6 with lane 1). We questioned whether these slow-migrating species were synthesized repair products (or expansions), or synthesis-free DNA structural changes. The production of the slower migrating species required HeLa cell extract, but was not a product of DNA synthesis, as they are only detected by Southern blot, not evident in native radio-incorporation reactions, and denaturing conditions only yielded the correct repair products (Supplementary Figures S3D and S3E). Thus, the slower-migrating species produced in the nick-in-repeat reactions are altered DNA structural conformations that have not incurred any synthesis. Considering that this conformational interconverstion of the nick-in repeat slip-outs required cell extracts, it is possible that SSBs, known to melt and reanneal unusual DNAs structures, may be involved in their resolution.

The striking difference between nick-in-repeat and nick-in-flank Δ(CAG)20 slip-outs was the efficiency of repair, such that the nick-in-flank slip-out had considerable levels of unrepaired substrate remaining following repair (Figure 2B lane 2) relative to its equivalent nick-in-repeat slip-out which was very efficiently repaired, leaving almost no unrepaired substrate (Figure 2C, lane 2). Repair levels were molar, being assessed by Southern blot. The complete correct repair of the nick-in-repeat substrates was further confirmed by the radionucleotide-incorporated repair reactions, which yielded only the expected sizes of the correct nick-directed repair products, in contrast to the nick-in-flank substrates (Supplementary Figures S3D and S3E). Quantitatively, nick-in-flank substrate had a 13% repair efficiency, in contrast to the nick-in-repeat substrates which had 81% repair efficiency (~6-fold greater; Figures 2B and 2C, compare lane 2 quantifications). Further, repair of a 3’ nick-in-repeat substrate was also higher than the nick-in-flank at ~30% (~2.5-fold greater; Figures 2B and 2D; compare lane 2 quantifications). We conclude that repair efficiency is highly affected by nick location, with the nick-in-repeat substrate being more efficiently correctly repaired than the nick-in-flank substrates, regardless of the repeat sequence.

### RPA and Alt-RPA enhance slipped-DNA conformational changes and repair

To test the role of RPA and Alt-RPA in *in vitro* repair, we prepared HeLa extracts depleted of RPA2 via siRNA (Supplementary Figure S3F). Depletion of RPA2 is expected to result in coordinate depletion of RPA1 and therefore the whole RPA or Alt-RPA complex, as we previously demonstrated^44^. HeLa cells do not express detectable levels of RPA4 (Supplementary Figure S3F)^44,47^. The diminished functionality of the RPA-depleted extracts and rescued functionality by supplementation of the purified human recombinant RPA/Alt-RPA proteins was verified by SV40 *in vitro* DNA replication assay, where the RPA-depleted extract was previously demonstrated to function only in the presence of added RPA and is not supported by added Alt-RPA (Supplementary Figure S3G)^43^.

Performing *in vitro* repair, we observed that the RPA-depleted extract was less effective, but not ablated, for its ability to correctly repair both nick-in-flank or nick-in-repeat slip-outs (Figure 2B-D, compare lanes 2 and 3 in each panel). Supplementation of the depleted extract with purified recombinant RPA and Alt-RPA enhanced repair efficiency for all substrates tested, with enhancement of repair being sensitive on nick location. Repair was enhanced by ~15% for the 3’ nick-in-flank substrate and ~20% for its equivalent 3’ nick-in-repeat substrate, and was enhanced to a lesser degree, ~5% for the 5’ nick-in-repeat substrate (nick on the same strand as the slip-out) (Figure 2B-D; compare lane 3 with lanes 4 and 5 in each panel). Similar observations were made for theshort slip-out and G-T mismatched substrates (Supplementary Figures S3H and S3I). This suggests that RPA and Alt-RPA enhance repair of large slip-outs, small slip-outs, and G-T mismatches, but are not essential for their repair. Notably, for nick-in-repeat substrates, RPA completely resolved, and Alt-RPA reduced, the slow-migrating products which are likely a DNA structural transition intermediate (Figures 2C and 2D; white arrowheads in lanes 4 and 5 relative to lanes 2 and 3). Our results therefore suggest that RPA or Alt-RPA can prevent/reduce the formation of, and/or promote the repair of, these structural intermediates.

Enhancement of slipped-CAG repair, and facilitation of structural intermediate transitions, are specific for the human RPA/Alt-RPA complexes, as neither purified *Saccharomyces cerevisiae* RPA (scRPA) or *E. coli* single-stranded DNA binding protein (bSSB) were able to enhance slip-out repair or facilitate structural transitions of any of the nick-in-repeat slipped-DNAs (Figure 2I and 2J). This suggests the primate-specific human RPA/Alt-RPA complexes are uniquely able to enhance slipped-DNA repair. A likely explanation is that the human RPA/Alt-RPA complexes mediate species-specific protein-protein and protein-DNA interactions which enhance slipped-CAG repair.

### High levels of Alt-RPA, but not RPA, inhibits slipped-DNA repair

Since HD and SCA1 patient brains exhibit upregulation of canonical RPA and Alt-RPA (~2-fold for RPA and between ~2-10-fold for Alt-RPA, with ratios of Alt-RPA to RPA being between 2:1 and 10:1; Figure 1 and Supplementary Figure S1), we next tested how elevated levels of RPA/Alt-RPA affect slipped-DNA repair. Since Alt-RPA exerts a dominant negative effect on the DNA replication activity of canonical RPA^43,44^, we hypothesized that a similar effect might also occur with slipped-DNA repair processing.

When equal amounts of purified RPA and Alt-RPA (1:1 ratio) were added to RPA-deficient cell extract, nick-in-repeat DNA structural transitions and repair efficiency remained unchanged from adding RPA or Alt-RPA alone (Supplementary Figure S3J). Strikingly however, increasing ratios of Alt-RPA to RPA (starting from a 2:1 ratio) yielded increasingly inhibited repair, with nearly completely inhibited repair observed at a ratio of 9:1 Alt-RPA to RPA (Figures 2E and 2F, compare lane 4 to lanes 5-7). Addition of the highest concentration of Alt-RPA in the absence of RPA also inhibited repair to the same degree (Figures 2E and 2F, compare lane 4 to lane 8). In contrast, no inhibition of slipped-DNA repair was observed with increasing levels of RPA relative to Alt-RPA (ratios ranging from 2:1 to 9:1 RPA to Alt-RPA) for either substrate (Figures 2G and 2H, lanes 4-8). These results demonstrate that while low levels of Alt-RPA can mildly enhance repair (Figure 2B-D, Supplementary Figure S3J), high levels of Alt-RPA, approximating upregulations observed in HD and SCA1 patient brains, inhibit slipped-DNA repair (Figure 2E and 2F). Interestingly, inhibition of repair by Alt-RPA was not limited to slipped-CAG DNAs, as increasing levels of Alt-RPA also inhibited G-T base-base mismatch repair, in contrast to the enhanced G-T mismatch repair elicited by increasing levels of RPA (Figure 2K). These results suggest that Alt-RPA competes with canonical RPA for single-stranded regions prior to or during DNA repair. This difference in activity between Alt-RPA and RPA could either be caused by differences in interactions with the slipped-DNA substrates (differences in their ability to bind and/or melt unusual DNA structures), interactions with other repair proteins, or both.

It is noteworthy that the production of the slower-migrating SI-DNA isomers from the faster-migrating nick-in-repeat substrates, was more evident in the absence of added RPA or Alt-RPA. The addition of increasing levels of RPA diminished its detection coincident with the efficient production correct repair products. In contrast, increasing levels of Alt-RPA yieled only the starting material, no structural transition products or repair products (Figures 2E-2H, compare white arrowhead in lane 3 to lanes 5-8). This suggests that structural transitions, likely a heterogenous random coil nick-in-repeat conformation to an ordered intra-strand hairpin conformation, are occurring before repair processing starts.

### Alt-RPA has altered binding, poorly unwinds slipped-DNAs, and inhibits FAN1 cleavage

First, we tested for DNA-binding differences between RPA and Alt-RPA to radiolabeled linearized nick-in-repeat SI-DNA3 and SI-DNA4 substrates by electrophoretic mobility shift assays. While RPA and Alt-RPA both formed protein-DNA complexes with the slipped-DNAs, the bound species formed were markedly different (Figure 3A, Supplementary Figure S4E). RPA binding yielded two distinct shifted protein-DNA complexes with increasing RPA concentration, suggesting that at least two molecules of RPA bound to the slipped-DNAs in slower migrating bands (Figure 3A, Supplementary Figure S4E, lane 4 in each panel). In contrast, Alt-RPA binding yielded only one protein-DNA complex at all concentrations (Figure 3A, Supplementary Figure S4E, lane 7 in each panel). Both RPA and Alt-RPA have a binding site size of 20-30 nucleotides^33,43^; this indicates that the substrates contain at least 20-30 nucleotides of ssDNA initially, but the length of this ssDNA stretch increases as more RPA is included in the reaction and multiple RPA-DNA complexes form.

**Figure 3:**
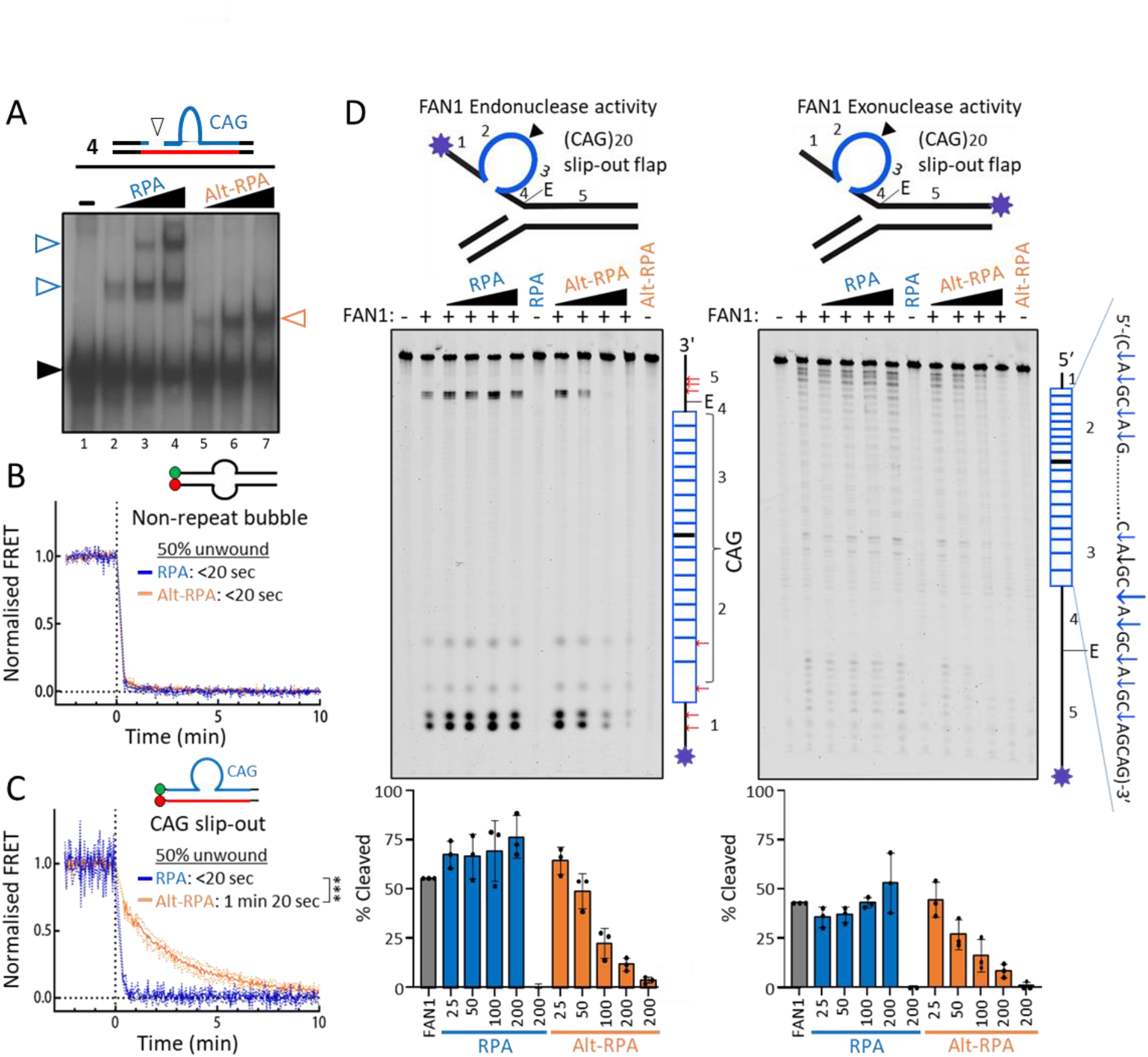
RPA and Alt-RPA bind slip-outs differentially and Alt-RPA cannot melt slip-outs, and Alt-RPA inhibits FAN1 endo- and exo-activity. A) DNA binding assays were carried out as described in Materials and Methods. Representative autoradiographs of RPA and Alt-RPA band-shifts using SI-DNA 3, SI-DNA 4, a linear DNA substrate containing 50 CTG/CAG repeats are shown, or a variety of single stranded, and unstructured or structured double stranded DNA Cy3 fluorescently labelled oligonucleotides. A) Non-radiolabelled DNA substrates (20 ng) were incubated with 600 ng (lanes 2, 5), 1200 ng (lanes 3, 6), or 2400 ng (lanes 4, 7) of RPA or Alt-RPA, as indicated, digested to release the repeat-containing fragment (EcoRI/HindIII), resolved on 4% native acrylamide, electrotransferred and probed. The positions of free DNA are indicated by black triangles; the positions of shifted protein-DNA bound complexes are indicated by white triangles hollow arrow head. B-C) Protein-mediated unwinding of DNA (melting) was observed by saturating the DNA with either protein complex, and observing the FRET signal over time. The time needed to reach 0.5 FRET (i.e. half of the DNA being unwound) was used to quantify the rate of substrate melting by each complex. D) FAN1 endo- and exo-nuclease activity is enhanced by RPA and inhibited by Alt-RPA. Purified FAN1 and RPA or Alt-RPA were incubated with FAM-labelled structured oligonucleotides (which mimic DNA overhangs that are generated during repair processing). DNA was resolved on a 4% urea denaturing gel and fluorescent signal visualized. A schematic on the right hand side of each gel outlines the labelled top strand (containing the CAG repeat) and where in the gel each fragment of DNA would be expected to migrate after electrophoresis. Red arrows outline exonucleolytic cleavage patterns elicited by FAN1, while blue arrows outline endonucleolytic cleavage patterns elicited by FAN1 within the CAG repeat. Nuclease activity was quantified densitometrically by comparing the intensity of the cleavage products to the intensity of the full length oligonucleotide band (top of the gel). N = 3 replicates.

Binding analysis to single-stranded, fully-paired duplexes, and various bubble DNA substrates revealed that Alt-RPA has lower affinity than RPA for ssDNA. With the different DNA substrates tested, initial shifts required ~2.5 to 3.5 higher levels, and saturation required ~2.5 to 5 higher levels, of Alt-RPA relative to RPA (Supplementary Figures S4A and S4B). We next probed for DNA unwinding differences between RPA and Alt-RPA, using a Förster resonance energy transfer (FRET) assay (schematic and calculation in Supplementary Figure S5D). First, we titrated purified RPA or Alt-RPA into different DNA substrates (duplex, bubble, CAG-slip-out, and hairpin) and calculated the equilibrium FRET value to determine the concentration of either complex needed to reach 50% DNA binding. Consistent with the band-shifts, we observed that more Alt-RPA is needed to reach 50% binding for all substrates (Supplementary Figure S4C). Next, we investigated the rates of RPA- and Alt-RPA-mediated DNA unwinding, by saturating the bubble or slip-out substrates with either purified RPA or Alt-RPA and monitoring the FRET values over time. Melting of the 8-nt bubble substrate was very rapid with no detectable difference observed between RPA and Alt-RPA - both reaching 50% unwound in <20 seconds (Figure 3B). Similar to the bubble-substrate, RPA rapidly unwound the non-repeat slip-out and CAG slip-out substrates in <20 seconds. In stark contrast, Alt-RPA was significantly slower at unwinding the CAG slip-out (1 minute 20 seconds, p<0.0017) and the non-repeat slip-out (1 minute 55 seconds, p<0.0004) substrates (Figure 3C, Supplementary Figure S4F). These data support that in addition to lower affinity binding to slip-outs, Alt-RPA also unwinds slip-outs at an exceedingly slow rate. Based on these results, slip-outs would be more biophysically stable in the presence of Alt-RPA as they are inefficiently bound and poorly melted by the Alt-RPA complex.

RPA’s ability to melt unusual DNA structures, including repeat-free hairpins, can modulate nuclease activities^35,74–76^. One hypothesis, based upon the differential ability of RPA and Alt-RPA to melt repeat slip-outs that we observed, is that these ssDNA binding complexes may differentially affect slip-out excision. FAN1 is a DNA nuclease identified as a potent disease modifier of seven CAG expansion disorders including HD and SCA1^12,14^. This link that has recently been supported to be by FAN1’s ability to regulate somatic CAG repeat instability in brains of HD mice through its nuclease activity. FAN1 cleaves slip-out DNAs with a unique specificity compared to non-repeat DNAs^62^. We assessed the effects of RPA and Alt-RPA upon the *endo*- and *exo*-nucleolytic activities of FAN1 on CAG-slip-out DNAs. RPA has a mild stimulatory effect upon *endo-* and *exo*-nulceloytic digestion by FAN1 on slipped-CAG DNAs (Figure 3D, left panel). In contrast, Alt-RPA strongly inhibited FAN1’s *endo-* and *exo*-nucleolytic digestion, preferentially blocking the cleavages in the repeat itself (Figure 3D, right panel). Consistent with the DNA binding differences between RPA and Alt-RPA, their ability to affect FAN1 digestion extended to a non-repeat unstructured flapped DNA (Supplementary Figure S4G). We conclude that Alt-RPA binds slipped-DNAs differently relative to RPA, Alt-RPA inhibits slip-out melting while RPA supports melting, and Alt-RPA inhibits FAN1 cleavage of slipped-DNAs while RPA promotes this activity. Seemingly, the ability and inability to melt slip-outs by RPA and Alt-RPA, respectively correlates with their respective abilities to enhance and inhibit FAN1 cleavage and slip-out repair.

### BioID reveals novel, unique and shared protein-protein associations for RPA/Alt-RPA

Functional overlap and distinctions between RPA and Alt-RPA, and the pathways in which these SSBs are involved, may be revealed by the proteins with which they associate. To generate proximal protein-protein associations we conducted BioID^77–80^ analysis for either RPA1, RPA2, RPA3, or RPA4 in living human cells (HEK293T). Proximal proteins were identified and processed as described^81–83^. For this pre-print, the BioID datasets have been abbreviated, with the raw data files being available through contact with the corresponding author (Dr. Christopher E Pearson). BioID reveals associations that are not necessarily limited to direct physical associations but can also include proteins that act in shared pathways in close proximity. The RPA1-4 BioID data are extensive. Below we focus upon interactions pertaining specficially to modifiers of CAG instability. Our RPA1-4 BioID results also identified subunit-protein associations with numerous HD/SCA disease modifiers, CAG/CTG disease pathogenesis, other repeat expansion diseases, chromatin biology, DNA metabolism, RNA metabolism, spontaneous DNA damage response in CAG/CTG disease tissues/models, and altered protein associations upon DNA damage. These are covered in the Supplementary Text.

Our experiments revealed previously characterized RPA interactors, including BLM, WRN, BRCA2, FANCJ, RAD50, MRE11, RAD51, NBN, ATRIP, P53, PCNA, polymerase II, BCAS2, RAD18, RECQL, SMARCAL1, UNG, etc (Supplementary Figure S5J)^84–86^. Previously reported Alt-RPA interactions with RFC, polymerases α and δ, and RAD51, were also detected in our BioID (Supplementary Figure S5J)^43,47^. While RPA and Alt-RPA do not physically interact with PCNA^43^, RPA indirectly associates with PCNA in multiple pathways - an association that was confirmed in our BioID (Figure 4A).

We also identified unique, shared, and novel associations (Figure 4A). In total we identified >2000 proximal protein associations for RPA1, RPA2, and RPA3, and ~700 associated proteins for RPA4, with a >5 log2-fold enrichment and p-value <0.01 relative to untransfected controls (Table 1, Supplementary Figure S5A-D^87^). Most proteins, 1674, were shared between RPA1, RPA2, and RPA3. Many, 581, were shared between all four subunits. Unique associations were identified for each subunit (60 to 150, ~4 to 8% of each subunit’s total interactions). That most associations were shared is consistent with RPA/Alt-RPA being predominantly heterotrimeric complexes. On the understanding that each subunit is necessarily part of either the canonical RPA (RPA1-RPA2-RPA3) or Alt-RPA (RPA1-RPA4-RPA3) heterotrimeric complex, unique interactions with RPA2 or RPA4 might reflect unique interactions with RPA and Alt-RPA, respectively, while unique interactions with RPA1 or RPA3 can be attributed to either complex. Gene Ontology (GO) term analysis of associations showed similar/proportional distributions of each RPA subunit for molecular function, biological processes, and cellular components, with minor differences being evident for RPA4 (Figure 4B, Supplementary Text)^88,89^.

### RPA upregulation inhibits somatic CAG repeat expansions in brains of SCA1 mice coincident with rescue of molecular, cellular and motor phenotypes

Towards assessing a possible *in vivo* effect of RPA upon somatic CAG expansions and hence disease, we overexpressed Rpa1 in the brains of mutant Ataxin-1 Q135 knock-in SCA1 mice, which were previously characterised^90^. We reiterate that mice do not have a functional *RPA4* gene (and therefore do not express Alt-RPA), hence upregulation of RPA1 leads to specific upregulation of canonical RPA complex, without inducing expression of Alt-RPA. This heterozygous mouse population was derived from the well-characterized Ataxin-1 Q154 heterozygous mouse model, which display ongoing tissue- and age-specific somatic CAG expansions^91–94^. The pattern of somatic CAG expansions in SCA1 and HD mice generally reflects that occurring in humans with SCA1 and HD, where CAG expansions were greater in the brain regions than peripheral tissues, with the striatum showing the largest and the cerebellum showing less, but not the lowest, CAG expansions in the CNS^3,95^. SCA1 mice, similar to SCA1 patients, exhibit gait and limb ataxia, dysarthria and dysmetria, and severe atrophy of the cerebellum and brainstem^93^. At the molecular level, vulnerable neurons demonstrate elevated levels of genome-wide DNA damage and ubiquitin-positive mutant expanded protein aggregates^93^.

Murine Rpa1 was overexpressed by injecting AAV-EGFP-Rpa1 into the subarachnoid space for broad brain delivery in 5-week-old. Previously, we demonstrated that AAV-Rpa1 overexpression rescued motor phenotypes (gait and rotarod), Purkinje neuron morphology, elevated levels of DNA damage in Purkinje neurons, and partially rescued impaired transcription, splicing, and abnormal cell cycle in these same SCA1 mice (Supplementary Figure S6A)^90^. It is noteworthy that a single injection of AAV-Rpa1 provided long-lasting recovery of motor function to Atxn1-KI mice for more than 50 weeks, with mice being assessed at 56 weeks of age.

Striatal AAV delivery in the SCA1 mice was confirmed by immunofluorescence of cytoplasmic EGFP in striatal DARPP-32-positive medium spiny neurons (MSNs) in comparison to no EGFP signal observed in non-injected SCA1 mice MSNs (Supplementary Figure S6B). We next confirmed that AAV-Rpa1 overexpression occurred within the striatum of the SCA1 mice, via ddPCR. We observed *Rpa1* RNA was upregulated (~2-fold, p=0.0057) in AAV-Rpa1 mice relative to AAV-EGFP mice and non-injected transgenic siblings (Figure 5A). *Rpa2* and *Rpa3* RNA levels were also upregulated in the same mice (Figure 5A), suggesting that Rpa1 expression mediates expression of Rpa2 and Rpa3, and that Rpa1 upregulation is sufficient to cause upregulation of the canonical RPA complex. This is consistent with previous reports where upregulation of human RPA1 prompts upregulation of RPA2 protein levels^96^. Notably, overexpression of *Rpa1* in mice will only affect the RPA complex, and not the Alt-RPA complex, as mice do not express RPA4. We then assessed for changes in CAG repeat expansion rates of AAV-EGFP-Rpa1 SCA1 mice relative to age- and inherited CAG length matched transgenic siblings overexpressing just EGFP by AAV-EGFP.

**Figure 5:**
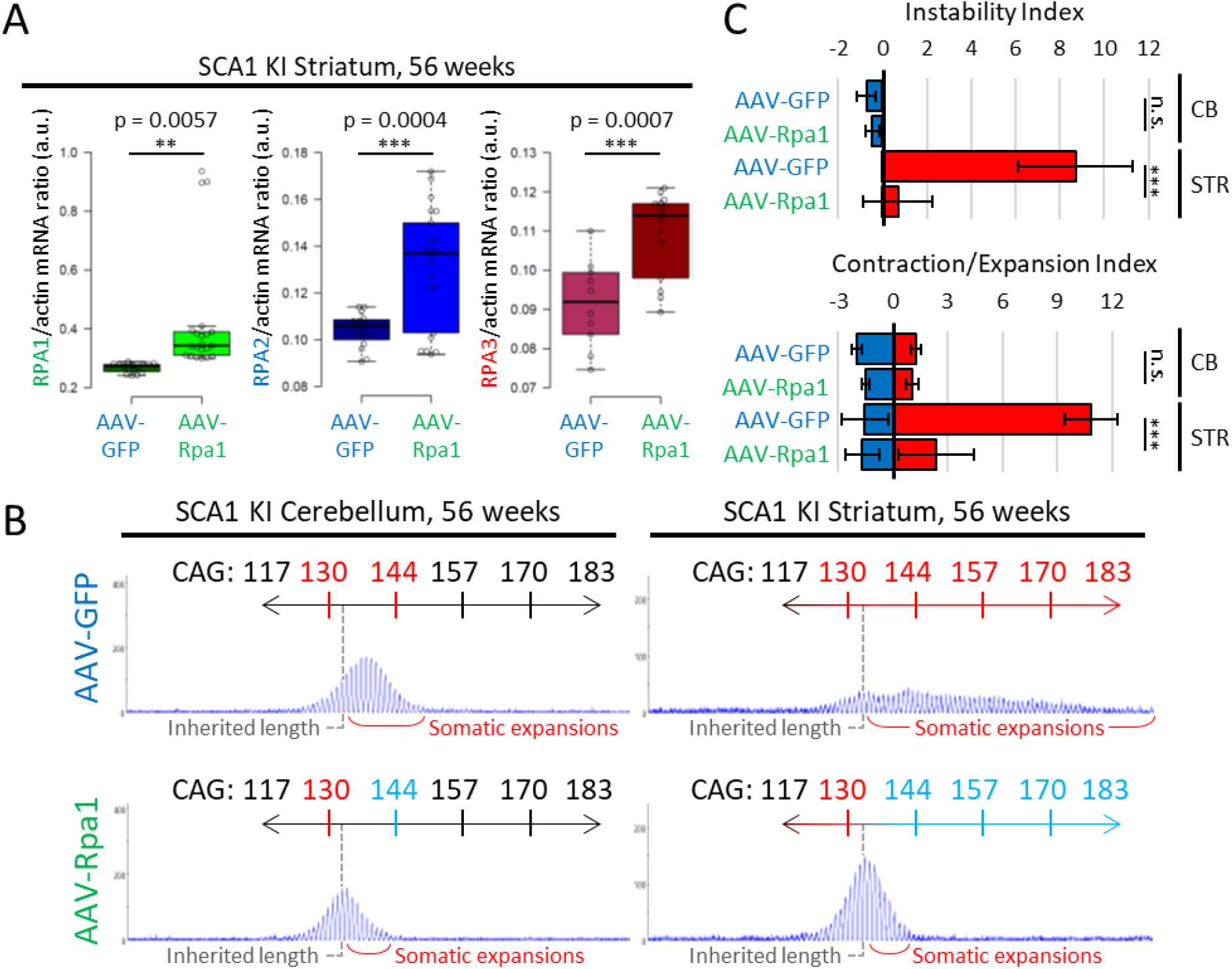
upregulation of canonical RPA in the striatum of a SCA1 mouse model inhibits somatic repeat instability. A) ddPCR results, graphed in box-plots, from the striatum of GFP- and Rpa1-overexpressing SCA1 mice reveals variable RNA upregulation of all subunits. n = 3 GFP overexpression mice, and 5 Rpa1 overexpression mice. Statistics: unpaired student t-test comparing means. B) Representative fragment length analysis scans of GFP- and Rpa1-overexpressing 56-week-old SCA1 mouse cerebellum and striatum. Gray bar designates the inherited repeat length, while red brackets label ongoing somatic expansions with age. C) Average instability and expansion/contraction indices for all GFP- and Rpa1-overexpressing SCA1 mouse cerebellum and striatum. *** = p < 0.001

Somatic CAG repeat expansions were then assessed via fragment length analysis in a series of mice all matched for inherited length of (CAG)~135. Within the cerebellum, we observed modest but consistent somatic CAG expansions from the inherited repeat length in AAV-EGFP SCA1 mice (Figure 5B, individual mice in Supplementary Figure S6B). This is consistent with previous reports in SCA1/HD mice and humans, where the bulk cerebellum shows low expansion levels relative to other brain regions, but is still actively expanding^3,94,95,97^. Upregulation of Rpa1 ablated somatic CAG expansions in the cerebellum (Figure 5B) - a trend consistent for each mouse (see individual mice in Supplementary Figure S6C). The lower levels of expansions in cerebellum did not allow its ablation by Rpa1 overexpression to be statistically significant (p=0.76).

The striatum of AAV-EGFP SCA1 mice exhibited extremely high levels of repeat expansions from the inherited length, with repeats ranging upwards of 193 CAG repeats - accruing >60 CAG repeats more than the inherited length of ~135 repeats (Figure 5B, individual mice in Supplementary Figure S6B). This is consistent with previous reports in striatum of SCA1/HD mice and humans^3,95,97^. Strikingly, AAV-Rpa1 upregulation within the SCA1 mouse striatum completely inhibited somatic repeat expansions, with the vast majority of repeat lengths clustering around the inherited repeat length of ~135 repeats (Figure 5B, individual mice in Supplementary Figure S6B). Degrees of repeat expansion were quantified by repeat instability indices, which calculate the average change in CAG repeat lengths relative to the most common repeat length (represented by the highest peak)^98^. The effect of Rpa was highly significant at reducing CAG expansions within the striatum (p=8.63e^−10^) (Figure 5C, individual mouse indices in Supplementary Figure S6C). Parsing the overall instability indices to contraction and expansion indices (Figure 5C) demonstrates that Rpa1 overexpression reduces the instability index by inhibiting somatic CAG expansions, rather than by inducing repeat repeat contractions.

### RPA upregulation reduces genome-wide DNA damage and mutant ATXN1 aggregation

Spontaneous elevated DDR markers are associated with several repeat diseases, including SCA1 and HD. These have been reported in patient tissues, cells, and mice^90,99–107^. Considering this and the extensive reports on the role of RPA in DDR pathways^108–111^, and our findings that the RPA subunits associate with numerous DDR proteins (Figure 4C), we assessed the effect of AAV-Rpa1 upregulation on DDR markers in treated SCA1 mouse brains. We observed that AAV-Rpa1 upregulation reduced γ-H2AX and 53BP1 (two markers of DNA DSBs) fluorescence intensity in cerebellar Purkinje neurons (Supplementary Figure S7A and S7B). We also observed significant reductions in both γ-H2AX (p=0.0005) and 53BP1 (p=0.0007) fluorescence intensity in DARPP-32-positive MSNs within the striatum of the AAV-Rpa1, relative to AAV-EGFP SCA1 mice (Figure 6A and 6B). This suggests that RPA upregulation can suppress spontaneous DDR, coincident with its ability to suppress somatic CAG expansions.

**Figure 6:**
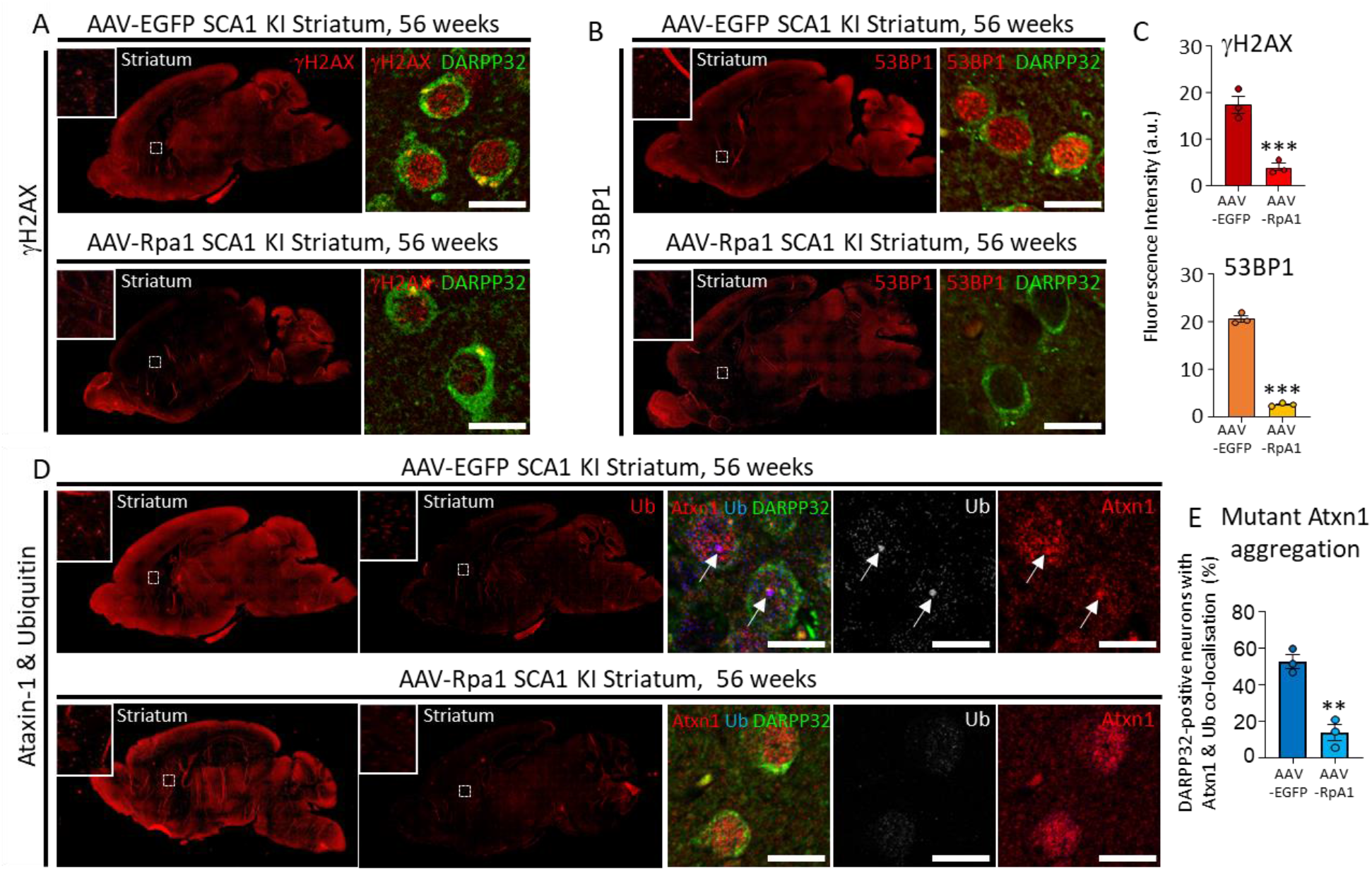
upregulation of canonical RPA in the striatum of a SCA1 mouse model reduces neuronal DNA damage and mutant Ataxin-1 aggregation. A-B) Representative confocal microscopy immunofluorescent images of γ-H2AX and 53BP1 signal in DARPP32-posititive striatal medium spiny neurons (MSNs) in GFP- and Rpa1-overexpressing SCA1 mice. C) quantification of fluorescent intensity of γ-H2AX and 53BP1 signal in GFP- and Rpa1-overexpressing SCA1 mouse striatal medium spiny neurons. n = 3 mic per group, 3 replicates with at least 30 neurons per replicate. *** = p < 0.001. D) Representative confocal microscopy immunofluorescent images of Ataxin-1 and ubiquitin co-staining within DARPP32-posititive striatal medium spiny neurons (MSNs) in GFP- and Rpa1-overexpressing SCA1 mice. E) quantification of the percentage of DARPP32-positive MSNs with ubiquitin-positive Atxn1 aggregation in GFP- and Rpa1-overexpressing SCA1 mouse striatal medium spiny neurons. n = 3 mic per group, 3 replicates with at least 50 neurons per replicate. ** = p < 0.01.

Aggregation of expanded polyglutamine proteins is a disease biomarker that is linked to polyQ size, and hence CAG repeat size^29,112–117^. Considering overexpression of RPA could suppress somatic CAG expansions at the DNA level, we postulated that this would extend to mutant ATXN1 polyQ lengths, thereby suppress mutant ATXN1 aggregation. To assess this, we analyzed the levels of mutant ATXN1 aggregates in DARPP-32 positive MSNs in AAV-Rpa1 relative to AAV-EGFP SCA1 mice (Figure 6D and 6E). We distinguished mutant ATXN1 aggregates from non-aggregated ATXN1 by double-staining for ATXN1 and ubiquitin, as aggregates form ubiquitin-positive nuclear inclusions which are resistant to proteasomal degradation^118^. While the AAV-EGFP SCA1 striatal MSNs showed a high degree of ataxin-1/ubiquitin double-positive inclusions (~52%), the number of double positive inclusions was significantly reduced in AAV-Rpa1 overexpressing SCA1 striata (~13%, p=0.003) (Figure 6E). This suggests that RPA-mediated suppression of somatic CAG expansions in striatal MSNs manifests in strong reductions of mutant ATXN1 polyQ aggregation.

## DISCUSSION

Repeat expansion mutations will involve single-stranded DNA intermediates and unusual DNA structures - requiring single-strand DNA binding protein complexes (SSBs) to stabilize, protect, melt, and anneal individual strands and recruit appropriate DNA repair proteins. Here we investigated two SSBs, RPA and Alt-RPA. RPA has been intensely documented (>3000 studies) to participate in every kind of DNA transaction. Alt-RPA is less understood, limited to 6 studies^42–47^. The data presented here support an Alt-RPA↔RPA antagonistic interaction, where Alt-RPA acts to oppose at least some of the functions of RPA.

We developed a working model for the role of RPA and Alt-RPA in CAG repeat instability (Figure 7). We find that RPA and, to a greater degree Alt-RPA, are upregulated in HD and SCA1 patient brain regions vulnerable to degeneration. RPA enhances correct repair of slipped-DNA (intermediates of expansion mutations). In contrast, high levels of Alt-RPA inhibit slip-out repair, where retention of the excess repeats causes expansions. Mechanistically, the differential repair outcomes by RPA/Alt-RPA can be explained by their differential abilities to bind, melt or retain slip-outs, and enhance or inhibit slip-out excision by FAN1, a nuclease which diminishes somatic expansions in HD mouse brains. Differential associations with proteins known to mediate somatic instability may also influence how RPA and Alt-RPA promote stability or instability, respectively. Enriched associations for RPA4 vs. RPA2, such as MSH3 and XPG with RPA4, may also influence how other identified proteins associate with RPA1 and RPA3 (components of both complexes). These data cumulatively support a model where RPA guards against somatic CAG expansions, thereby diminishing severe phenotypes, while Alt-RPA blocks RPA’s activity, promoting expansions and worsened phenotypes (Figure 7). Supporting this model, overexpression of RPA in SCA1 mice prevents somatic expansions in the brain, diminishes molecular markers of disease (DDR and polyQ aggregates), and rescued neuron morphology and motor phenotypes. These findings reveal RPA and Alt-RPA as active players in CAG repeat stability and instability, respectively. In bacterial and yeast models of CTG and GAA repeat instability, where repeat contractions predominate, an absence of the single-strand binding proteins, SSB and RFA, led to enhanced repeat contractions^40,41^ – consistent with our findings that metazoan RPA is required to protect against the predominating repeat expansions.

**Figure 7:**
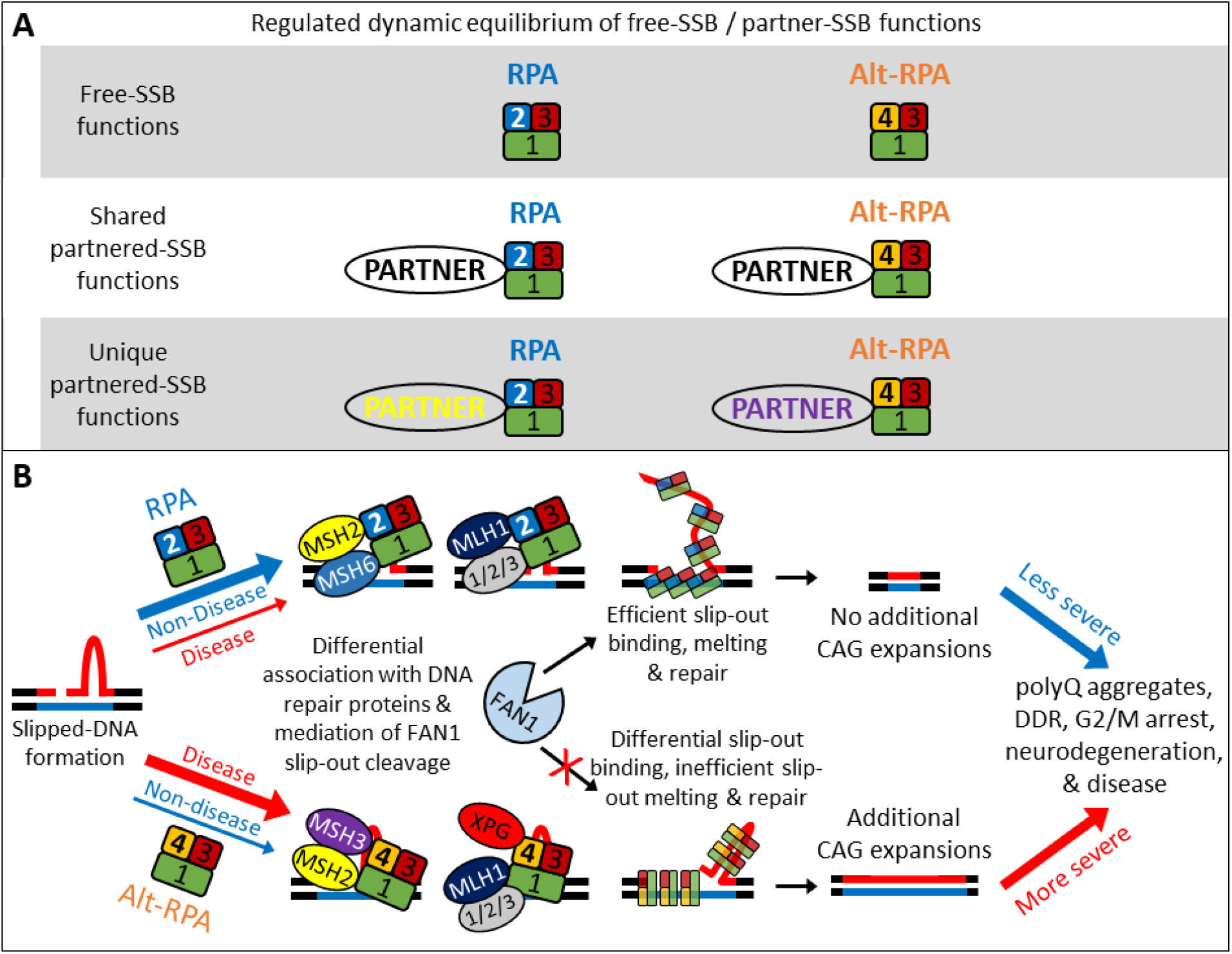
Working model for RPA and Alt-RPA on CAG/CTG expansions. Working model for the role of RPA and Alt-RA on CAG repeat instability and pathogenesis outlined that RPA and Alt-RPA are likely eliciting opposing roles on CAG repeat instability and pathogenesis. At slipped CAG structures, RPA is likely enhancing repair by efficiently melting slip-outs, thereby leading to no CAG expansions and less severe disease presentation, while Alt-RPA inefficiently melts slip-outs and thereby inhibits repair. Differential repair outcomes are also likely resulting from differential associations with DNA repair proteins known to modulate repeat instability, as well as differential effects on the activity of these proteins at the repeat (including antagonistic roles for RPA and Alt-RPA in promoting or inhibiting, respectively, FAN1 nuclease activity at slipped-CAG structures). We hypothesize that these antagonistic roles result in RPA inhibiting, and Alt-RPA promoting, somatic CAG expansions and pathogenesis.

Our BioID protein-interactomes, representing the first unbiased interactomes for RPA and Alt-RPA subunits, provides a foundation for the pathways involved in repeat metabolism in disease and non-diseased states. Unique associations of RPA and Alt-RPA with proteins known to regulate somatic expansions could be insightful. For example, the MSH2-MSH3 and MLH1-PMS2 complexes, are required to drive CAG expansions in brains of various HD mouse models, and apart from MSH2, each has been identified as disease modifiers in human HD and several SCAs. In contrast, MSH6 has been associated with stabilizing CAG tracts^119–127^. Differential associations with these proteins could have substantial effects on the rates of somatic expansions. For example, since RPA4 but not RPA1, RPA2, or RPA3, preferentially associates with MSH3 (Figure 4C), one can imagine Alt-RPA working with MutSβ, (MSH2-MSH3) to mediate CAG repeat expansions, possibly through retention of the excess repeats in poorly repaired slip-outs (Figure 2). Our data now reveal RPA1-4 as key players in CAG instability, despite the fact that RPA1-4, like MSH2, were not identified in the disease modifier screens^11–15,128^. Beyond repeat diseases, these RPA subunit BioID data provide a myriad of potential Alt-RPA↔RPA interactions which can shed light on a many metabolic pathways.

Evidence of the effects of altered RPA subunit expression offer insight to the extent of the Alt-RPA↔RPA antagonistic interaction. RPA is essential for viability^129^, and dysregulation of its expression levels can be clinically and molecularly impactful. Increases or decreases in RPA expression has distinct effects. In humans, modest increases (~1.5-fold) or decreases (~0.8-fold) in canonical RPA levels, via chromosomal microduplications or microdeletions of *RPA1* (among other genes), is associated with 17p13.3 microduplication/microdeletion disease syndromes^130–132^. Haploinsufficiency of *RPA1* in humans causes defective ATR-dependent DDR and G2/M checkpoint arrest, which is rescued upon exogenous RPA1 expression^130^. Experimental depletion of *RPA1* causes most cells to arrest in G2/M and subsequently die^44,133^. Duplications of *RPA1* lead to altered DDR and cell cycle changes, both distinct from RPA haploinsufficiency^131^.

Experimental over-expression of either RPA1, RPA2, or RPA3 subunits in human cells leads to endoreduplication (re-initiation of chromosomal replication prior to nuclear division leading to enriched tetraploidy) and attenuated DSBs and chromosomal instability^131,133^. This supports a direct link of RPA overexpression in suppressing elevated levels of γ-H2AX and 53BP1 observed in the SCA1 mouse brains (Figure 6, Supplementary Figure S7)^90,99–107^. Overexpression of RPA4 in absence of exogenous stress also leads to accumulation of DSBs (γ-H2AX foci), and subsequent cell death^44^. Thus, over-expression of RPA4 in unstressed cells induces a DDR, similar to the spontaneously activated DDR in SCA and HD patient cells and brains. That both depletion of canonical RPA and overexpression of Alt-RPA lead to DNA damage and apoptosis^131,133^, further supports an Alt-RPA↔RPA antagonistic interaction. Overexpression of RPA4 in SCA and HD patient cells and brains may be related to the accumulation of DSBs (γ-H2AX foci), re-entry of neurons into the cell-cycle, G2/M arrest, and subsequent cell death (neurodegeneration) occurring in HD patient tissues and HD mice^134,135^. While such a link is possible, whether the levels of HD-related over expression of RPA and Alt-RPA expression change are causally related to neurodegerneartion cannot, at this time, be definitively linked.

Increased and decreased levels of RP2 and RPA4, respectively, have been observed in many cancers, supporting Alt-RPA↔RPA antagonistic roles in chromosomal replication and proliferation^47,136^. Humans inheriting Xq21.33 duplications encompassing only the *RPA4* and *DIAPH2* genes are phenotypically normal (males and females), suggesting that Alt-RPA dosage variations may not be pathogenic^137^. However, somatically elevated RPA4 levels are associated with better prognosis in gastric cancers, suggesting that RPA4 levels can be important in disease states^136^. It is tempting to speculate that an antagonistic role of Alt-RPA upon DNA replication/proliferation may be linked to the reduced cancer incidence in HD patients^138–142^. It is also possible that Alt-RPA contributes to the greatly diminished pool of proliferating neuroprogenitor cells in human HD fetal brains and the depletion of postnatally generated striatal neurons in adult HD patients at advanced disease stages^135,143^.

To test the *in vivo* effects of RPA upregulation we used a SCA1 mouse model that faithfully replicates numerous pathological features of human SCA1 disease^93^. The ablation of rampant CAG expansions in the SCA1 mouse striatum by RPA overexpression coincides with reduced disease phenotypes. It seems reasonable to consider these phenotypic benefits were a direct result of minimizing progressive somatic CAG expansions. This strengthens the concept that therapeutic targeting of somatic CAG expansions will be phenotypically beneficial. Determining the *in vivo* mechanism by which Alt-RPA might regulate somatic expansions will be challenging. Analysis of RPA4 overexpression in mice is complicated by mice having a non-functional, non-expressed *Rpa4* pseudogene. As such, the *in vivo* role that RPA4 might be playing in CAG expansions or disease pathogenesis cannot currently be tested.

Patterns of instability between different brain regions is similar, but not identical, amongst most CAG/CTG expansion diseases (HD, SCA1, SCA3/MJD, DRPLA, and DM1), and are reflected in the respective mouse models. Typically, larger somatic expansions are present in the striatum and smaller expansions arise in the cerebellum. In HD, the potential contribution of somatic expansions to disease is likely, as the largest CAG expansions in the striatum directly correlates with the brain region most vulnerable to rampant and early degeneration^144^. Moreover, the levels of somatic expansions correlates with disease age-of-onset, further supporting a causal relationship of expansions to disease^4^. In contrast, in SCA1 the potential contribution of somatic expansions to disease has received limited attention, as a causal relationship between the regional degrees of CAG expansions and neurodegeneration was not obvious in post-mortem SCA1 brains. Specifically, the cerebellum, vulnerable to severe degeneration early in disease, has smaller expansions relative to the large expansions in the striatum which is relatively spared from degeneration in SCA1^145–150^. However, an apparent absence of correlation between instability and disease relies upon disease being solely due to neurodegeneration, which it is not. For example, HD and SCA1 patients can be clinically affected prior to any observed neurodegeneration^151–156^. Recent findings reveal that while the cerebellum degenerates early in SCA1, the striatum does degenerate later and more rapidly (with detectable changes in less than a year) later in the disease course. Notably, MRI-derived volumetric decreases of the striatumcorrelate with disease progression (decline in motor function) in SCA1^52,53^. Thus, neurodegeneration does not explicitely dictate disease presentation, as cellular dysfunction in the absence of neurodegeneration can also mediate pathogenesis. Our data supports that the striatum, and somatic repeat expansions within them, are contributing to SCA1 mouse pathogenesis. RPA overexpression correlating with reduced disease phenotypes suggests that inhibition of somatic expansions can be beneficial for allaying disease in SCA1. Therefore, somatic CAG expansion may drive the rate of various aspects of disease pathogenesis in many different cell types in both HD and in SCA1, regardless of a direct coincidence with levels of neurodegeneration.

A contribution of somatic CAG expansions to HD disease is likely, where the largest expansions occur in brain regions most vulnerable to degenerations. Moreover, screens for modifiers of HD onset or progression have revealed multiple DNA repair genes known to be involved in somatic CAG expansions. A contribution of somatic CAG expansions to SCA disease is unknown, but the similarity of DNA repair genes identified in disease modifier screens for multiple SCA, supports this suspicion^12^. Both HD and SCA1 show high somatic CAG expansions in the cortex and in striatum, regions that are the most vulnerable to degeneration in HD, but less so in SCA1. In SCA1 the cerebellum and brainstem are most vulnerable to degeneration, particularly losing the cerebellar Purkinje cells. It is clear that neurodegeneration of the cerebellum is important to disease in SCA1, however, degeneration in the cerebellum does not account for all SCA1 patient or mouse model phenotypes^52,157,158^. We see mild CAG expansions in the SCA1 murine cerebellum, likely missed in SCA1 humans due to the massive cerebellar cell loss that they, but not mice, incur. We also see rampant expansions in the striatum. During SCA1 disease progression, pathology extends beyond the cerebellum and brainstem to involve the striatum and temporal lobe^52,157,158^. Similarly, during HD progression pathology extends beyond the striatum to involve cerebellar Purkinje cells, brainstem nuclei, and peripheral tissues^54,159–164^. Disease pathogenesis may arise from affects in different brain regions and the relative contribution of instability. That over-expression of RPA, a previously unconceived modifier, can diminish expapsions in brains of SCA1 mice, and that this diminished some disease pehenotypes in the mice, supports the concept that somatic CAG expansion may also drive SCA1 disease.

In summary, we reveal new functions of canonical RPA and of the understudied Alt-RPA - highlighting its antagonistic effects upon RPA functions. Questions and implications stemming from our work include i) might Alt-RPA RPA antagonistic interactions impact RPA function in the many other DNA repair processes that require RPA? ii) Might regulated Alt-RPA RPA interactions sustain cells in a post-mitotic state or ensure high-fidelity DNA repair processes? Indeed, perturbation of the relative concentrations of RPA4 and RPA2 in disease or cancerous states may disrupt the Alt-RPA-RPA interaction equilibrium. This implies that most forms of DNA metabolism in primates could be affected by Alt-RPA.

## MATERIALS & METHODS

### Patient tissue sample collection, preparation, and patient descriptions

Post-mortem patient tissues were provided by the Neurological Foundation Human Brain Bank with institutional ethics approval #011654 (7 HD patients and 7 unaffected individuals; striatum, cerebellum, frontal pole) directed by RLMF and MAC. ARLS provided 3 HD patients and 3 unaffected individuals; striatum and cerebellum, and the National Ataxia Foundation Biobank (3 SCA1 patients; cerebellum). Tissues were collected from patients using previously characterised protocols^165^. Briefly, unfixed brain is sectioned into discrete blocks and frozen with powdered dry ice, double wrapped in aluminum, and then stored at −80°C until processing. Known clinical information is outlined below in Table 1.

### zQ175 HD mouse model description, handling, and tissue collection

Animal protocols were approved by the Children’s Hospital of Philadelphia Institutional Animal Care and Use Committee. Heterozygous zQ175 mice and littermate controls were housed on a 12-hour light/dark cycle in a temperature- and humidity-controlled environment with *ad libitum* access to food and water. Mice were anesthetized with ketamine/xylazine and perfused with ice-cold saline. Striatum and cerebellum samples were immediately collected on ice, flash frozen in liquid nitrogen, and stored at −80°C.

### SCA1 KI mouse model description, handling, and tissue collection

Handling, ethics, and tissue harvesting was previously described^90^. All mouse experiments, handling, and sacrifice were performed in strict accordance with the Guidelines for Proper Conduct of Animal Experiments by the Science Council of Japan. Mice were euthanized with ethyl ether, and tissues were collected within 5 minutes of death. Tissues were immediately frozen in liquid nitrogen and then kept in −80°C until processing. Mutant Atxn1 knock-in mice were crossed with background mice (C57BL/6J) during breeding. After multiple crosses, heterozygous knock-in mice with 125-140 repeats were used for all subsequent experiments, with non-transgenic siblings being used as controls.

### Patient derived cell line culturing

Q43, Q40, and control 1 cell lines were a gift from Dr. Ray Truant (McMaster University) and were previously characterised^166^. Some lines were purchased from the Coriell Biorepository; Q43 HD line (code: GM02191), Q45 SCA1 line (code: GM06927), Q53 SCA3 line (code: GM06153). Control cell line 2 and 3 were a gift from Dr. Guy Rouleau (McGill University) and control cell line 4 and 5 were a gift from Dr. Elise Heon (University of Toronto; C4 and C5). For long-term storage, cells were immersed in Cellbanker 1 (Amsbio, catalogue #11888) and frozen in liquid nitrogen. All cells were cultured in DMEM (10% FBS, 1% supplemented L-glutamine, 1% Penicillin-Streptomycin) at 37°C with 5% CO2. Cells were plated at ~50% confluency, and split at 85-95% confluency by Trypsin. Viability was checked using Trypan Blue exclusion tests during all splits and prior to experimentation; a viability of 90% or greater was maintained for cells prior to experimentation.

### RNA preparation from patient and mouse brain tissues

Tissues (stored and −80°C and kept immersed liquid nitrogen during handling) were crushed with a frozen metal mortar and pestle partway buried in dry ice, and frozen crushed tissues were immediately transferred to a 1.4 mm Acid Washed tube pre-filled with Zirconium Beads and 300-1000 μL of TRIzol reagent. Smaller tissues were directly inserted into tubes without crushing. Tubes were inverted to ensure immersion of the whole tissue, and was placed at room temperature for 10 minutes to allow the TRIzol reagent to denature and remove proteins bound to RNA. Tubes were placed on ice after ten minutes and then placed in a MagNA Lyser Instrument (Roche; item #03358968001). Tubes were oscillated at 7000 OSC 3 times for 20 seconds each oscillation, with a 3-minute incubation on ice between each 20 second oscillation. TRIzol was transferred to a different tube, RNA precipitated by an equal volume of 100% EtOH and then purified using the Direct-zol RNA purification kit using the manufacturers protocol, which includes in-column DNase treatment (Zymo research; catalog # R2071). Whole RNA was reverse transcribed using the SuperScript IV First-Strand Synthesis System kit using the manufacturers protocol (ThermoFisher Scientific; catalogue #18091050).

### Droplet digital PCR (ddPCR) for RNA transcript expression quantification

FAM and HEX fluorophore labelled probes specific for each RNA target of interest was ordered from Bio-Rad (pre-made probe designs), and manufacturer’s “PrimcePCR ddPCR gene expression probe assays” protocol was used for ddPCR reactions. In brief: 10-50 ng of total cDNA (depending on target abundance, empirically derived from preliminary runs) was mixed with: 1) 2x ddPCR Supermix for Probes (no dUTP), 2) 20x target primers/probe mix (FAM), and 3) 20x reference primers/probe (HEX), topped up to a final reaction volume of 20 μL with DNase-/RNase-free water. Plate was sealed with aluminum, mixed well and centrifuged briefly to collect the reaction, and then kept at room temperature for 3 minutes to equilibrate the reaction temperature to room temperature. Droplets were generated using the Automated Droplet Generator (Bio-Rad; catalogue number: 10043138). Plate containing generated droplets was re-sealed with aluminum, and then subjected to PCR in a C1000 Thermal Cycler (Catalog #185-1197) to the following cycles: 1) 1x 95°C, 10 minutes, 2) 40x (94°C, 30 seconds followed by 55°C, 1 minute), 3) 1x (98°C, 10 minutes), 4) held at 4°C until further processing. A ramp rate of 2 °C/second was used for all the cycles. After cycling, the plate was transferred to a QX200 Droplet Reader (Bio-Rad; catalogue #186-4101) for fluorescent detection, and was analysed using the Bio-Rad QuantaSoft Software. All experiments were conducted using at least 3 technical replicates.

### Quantitative reverse transcription PCR (qRT-PCR)

RNA isolation from patient brain tissues, and cDNA generation performed as described in the ddPCR methods above. mRNA quantification was performed using TaqMan probes (Life Technologies) and TaqMan Universal PCR Mix (ThermoFisher Scientific; catalogue # 4304437) on a 7500 Real Time PCR System (Applied Biosystems). Gene expression was normalized to 18S rRNA. Delta CT values were calculated as Cttarget - Ct18S. Note: all experiments were conducted using at least 3 technical replicates.

### Protein lysate preparation (cultured cells)

RIPA Lysis and Extraction Buffer (ThermoFisher Scientific; catalogue #89901) was mixed with and appropriate volume of 100x Halt Protease and Phosphatase Inhibitor Cocktail (ThermoFisher Scientific; catalogue #78430) and kept on ice. Fibroblast cells were kept adhered to the culture flask, media removed, and washed twice with 1x sterile PBS. After the last wash, as much excess PBS was removed as possible and 150 μL - 700 μL of RIPA with protease inhibitor (depending on cell numbers) was added directly to the culture flask. Flask was tilted to allow RIPA to cover the whole surface that cells were grown on, and then placed flat (to ensure whole surface was covered with RIPA) on ice for 1 hour. Cells were then scraped on ice using a rubber scraper, and RIPA was collected into an epi-tube. Cells were then sonicated on ice using a microtip (amplitude 20, 15 cycles, each consisting of 1 second on and 1 second off), with the microtip being cleaned twice with water and then 70% ethanol (wiped dry with a kim-wipe) between each sample. Cell debris was pelleted by centrifuging at 21,000xg for 15 minutes at 4°C, and the supernatant collected for subsequent experiments. Supernatant was aliquoted into 100 μL aliquots to avoid repeat freeze-thaws and then stored at − 80°C between uses.

### Protein lysate preparation (tissues)

Tissues (stored and −80°C and kept immersed liquid nitrogen during handling) were crushed with a frozen metal mortar and pestle partway buried in dry ice, and frozen crushed tissues were immediately transferred to a 1.4 mm Acid Washed tube pre-filled with Zirconium Beads and 300-1000 μL of RIPA without detergent (50 mM Tris HCl pH 7.4, 150 mM NaCl, 1 mM EDTA) with an appropriate volume of 100x Halt Protease and Phosphatase Inhibitor Cocktail (ThermoFisher Scientific; catalogue #78430), on ice. Smaller tissues were directly inserted into tubes without crushing. Tubes were inverted to ensure immersion of the whole tissue, and then placed back on ice before processing in a MagNA Lyser Instrument (Roche; item #03358968001). Tubes were oscillated at 7000 OSC 3 times for 20 seconds each oscillation, with a 3-minute incubation on ice between each 20 second oscillation. To ensure complete tissue homogenisation, samples were briefly spun to assess the level of unhomogenized tissue left (if any), and additional oscillation cycles were performed as needed. An equal volume of RIPA double detergent (2%DOC, 2% Igepal, 2% Triton X-100) with with an appropriate volume of 100x Halt Protease and Phosphatase Inhibitor Cocktail (ThermoFisher Scientific; catalogue #78430) was added to each tube. Parafilm and clips were attached to the lid of each tube, and then incubated on a sample roller overnight at 4°C. The next day, tubes were centrifuged at 14000xg for 15 minutes at 4°C, and the supernatant collected for subsequent experiments. Supernatant was aliquoted into 100 μL aliquots to avoid repeat freeze-thaws and then stored at −80°C between uses.

### SDS-PAGE and western blotting

10 μg - 100 μg of protein lysate was used per sample (depending on target protein abundance, and kept as consistent as possible between samples on the same gel). Samples were prepared using 4x NuPAGE LDS Sample Buffer (ThermoFisher Scientific; catalogue #NP0007) and 10x NuPAGE Sample Reducing Agent (ThermoFisher Scientific; catalogue #NP0004), and denaturing the samples at 70°C for 10 minutes. The denatured samples were electrophoresed at 100-120 volts for 1.5-2.5 hours on NuPAGE 4-12% Bis-Tris Proteins Gels (ThermoFisher Scientific; catalogue # NP0321BOX) in NuPAGE MES SDS Running Buffer (ThermoFisher Scientific; catalogue #NP0002). Samples were run in parallel with Full range rainbow MW marker (ThermoFisher Scientific, catalogue #RPN800E) and/or HiMark Pre-stained Protein Standard (ThermoFisher Scientific, catalogue #LC5699).

Gels were wet-tank transferred to PVDF Western Blotting Membranes (Sigma-Aldrich, Cat #3010040001; activated in 100% methanol for 1-2 minutes prior to use) in tris-glycine (with 10-20% methanol) overnight (16-24 hours typically) at 4°C using a constant voltage of 20-30V. The next day, membranes were blocked in 5-10% w/v milk dissolved in 1xTBS + 0.1% Tween-20 (TBST) for 1 hour at room temperature. Blots are then incubated with primary antibody at room temperature for 2 hours using the same solution used for, washed 3 times in 1xTBST at room temperature (10 minutes/wash), incubated with secondary antibody at room temperature for 1 hour in the same solution used for blocking, washed 3 times in TBST at room temperature (10 minutes/wash), and then detected with ECL (GE Healthcare Amersham ECL™ Prime Western Blotting Detection Reagent, Cat #RPN2232) by autoradiograph. Densitometric quantification of bands is performed using Image Studio Lite Version 5.2 (LI-COR Biosciences).

### Antibodies used for western blotting

*Primary antibodies:* Anti-RPA2 cone 9H8 (1:1000, monoclonal mouse, Abcam catalogue #ab2175), Anti-RPA4 (1:4000-1:8000, sheep serum, home made), Anti-Actin Protein Antibody (1:30,000, mouse) (BD Transduction Laboratory, catalogue #612657). *Secondary antibodies:* Peroxidase-AffiniPure Sheep Anti-Mouse IgG H+L (1:2000, Cedarlane Labs, catalogue #515035062), Sheep IgG (H+L) Highly Cross-Adsorbed Donkey anti-Ovine HRP (1:2000-1:6000, ThermoFisher Scientific, catalogue #A16047)

### Antibodies used for IF

*Primary antibodies:* anti-phospho-H2AX (γ-H2AX) clone JBW301 (1:200, monoclonal mouse, Millipore Sigma catalogue #05-636), anti-53BP1 (1:5000, polyclonal rabbit, Novus bio catalogue #NB100-304SS), anti-Ataxin1 clone N76/8 (1:100, monoclonal mouse, EMD Millipore catalogue #MABN37), anti-ubiquitin clone P4D1 (1:1000, monoclonal mouse, Cell Signalling Technology catalogue #3936S), anti-DARPP32 clone 19A3 (1:200, monoclonal rabbit, Cell Signalling Technology catalogue # 2306), anti-calbindin clone EG-20 (1:2000, polyclonal rabbit, catalogue EMD Millipore #05-636), anti-calbindin clone CB-955 (1:2000, monoclonal mouse, Millipore Sigma catalogue #C9848). *Secondary antibodies:* Goat anti-Mouse IgG (H+L) Superclonal Recombinant Secondary Antibody Alexa Fluor 555 (1:200, ThermoFisher Scientific catalogue #A28180), Goat Anti-Rabbit IgG H&L Alexa Fluor 488 (1:200, abcam catalogue #ab150077), Goat anti-Rabbit IgG (H+L) Cross-Adsorbed Secondary Antibody Alexa Fluor 568 (1:200, ThermoFisher Scientific catalogue #A-11011), Goat Anti-Mouse IgG H&L Alexa Fluor 488 preadsorbed (1:200, Abcam catalogue #ab150117).

### Functional cell extract preparation

Cells are grown in 20 cm plates to ~70-80% confluence. Media is removed and cells are washed twice with ice cold hypotonic buffer (20 mM Hepes-KOH pH 7.8, 5 mM KCl, 0.15 mM MgCl2 and 0.1 mM DTT). Remove as much excess hypotonic buffer from washes as possible, add 300 μL of hypotonic solution with 3 μL of 100x Halt Protease and Phosphatase Inhibitor Cocktail (ThermoFisher Scientific; catalogue #78430), and then scrape the cells using a rubber scraper. Collect cells into a Dounce homogeniser. Dounce cells ~10-15 times using a tightly fitting pestle (B pestle). Homogenisation of cells can be checked on a slide under a light microscope, and additional homogenisation can be conducted as needed. Transfer extract to a larger volume conical centrifuge tube (15 mL or 50 mL) and let stand on ice for 30 minutes. Centrifuge at 1700xg for 10 minutes at 4°C to pellet large cell debris, then transfer to high-speed centrifuge tubes and centrifuge again at 12,000xg for 10 minutes at 4°C to clarify the extract further. Remove supernatant and freeze as beads by dripping into liquid nitrogen, and then store in −80°C.

### *In vitro* repair reaction and Southern blotting

Substrate generation and repair reactions were performed as previously described^63,64^. In brief, each repair reaction consists of 1 μL slipped-DNA substrate, ATP, rNTP-ATP, calf phosphatase, calf kinase, dATP, dCTP, dGTP, dTTP, cell extract, with and without supplementation with purified protein. The reaction is incubated at 37°C for 1 hour, after which the reaction is stopped by 2% SDS, 2 mg/mL proteinase K, 0.05 M EDTA and incubating for another hour at 37°C. Phenol:chloroform extraction is performed and then DNA purified using MinElute Reaction Cleanup Kit according to the manufacturer’s protocol (Qiagen, catalogue # 28206). The DNA is then digested with *Eco*RI and *Hin*dIII overnight at 37°C. The next day the DNA is electrophoresed at 200 volts on a 4% polyacrylamide gel in 1x TBE for 1 hour and 25 minutes. The electrophoresed DNA is then transferred from the gel to a nitrocellulose membrane using a Owl HEP Series Semidry Electroblotting System (ThermoFisher Scientific, catalogue #HEP-1). The transferred membrane is then immersed in denaturing solution (1.6% w/v NaOH pellets in ddH2O) for 20 minutes at room temperature with gentle agitation, renatured in Southern neutralising solution for 20 minutes at room temperature with gentle agitation, and then washed in 5x SSPE for 20 minutes at room temperature with gentle agitation. Following this, the membrane is rolled into a glass hybridization tube and blocked with salmon sperm DNA in Southern prehybridization solution for an hour at 42°C. After this, a 32P radioactively labelled probe complimentary to the DNA is added to the prehybridization solution and allowed to hybridize overnight at 42°C. The next day, the radioactive probe in prehybridization solution is removed and the blot is washed 3x with Southern wash solution (0.1% SDS in 0.1% SSPE v/v in water) - each wash being at least 30 minutes long at 65°C). Lastly, the membrane is exposed to an autoradiograph to visualise the DNA.

### siRNA administration to cultured cells

The RPA2 siRNA was from Santa Cruz biotechnology (catalogue #sc-38230). siRNA was used according to the manufacturer’s specific protocols.

### *In silico* analysis of canonical-RPA and Alt-RPA

Human RPA2 and RPA4 sequences were extracted from Uniprot and aligned using Clustal Omega. The RPA4 structure model and associated model prediction data were downloaded from AlphaFold (accession Q13156) and analysed using Pymol and APBS.

### Electrophoretic Mobility Shift Assay (EMSA; aka band-shift assay)

Radioactively or fluorescently labelled DNA and proteins/compound are incubated at room temperature for 15-30 minutes in a reaction containing purified proteins and DNA in a binding buffer (3mM Hepes pH 7.9, 16 mM NaCl, 0.04 mM EDTA, 0.2 mM DTT, 0.06 mg/ml BSA, and 2% glycerol). Following this incubation, 1-2 μL of 10x native sample binding dye (50% glycerol with Bromophenol Blue) are added to the reaction and loaded onto a 4-8% polyacrylamide gel or 1-1.5% agarose gel as quickly as possible. The reaction is electrophoresed for 1-3 hours at room temperature or overnight at 4°C. For fluorescently labelled DNA, gels are visualised using a fluorescent detection system (typically an Amersham Typhoon laser-scanning platform or Bio-Rad ChemiDoc MP imaging system). For radioactively labelled DNA, gels are dried to a Whatman paper and then exposed to autoradiograph to detect band shifts.

### Kinetic FRET binding and melting assays

Two complimentary oligonucleotides with a Cy3 or Cy5 fluorophore were annealed to one another such that the fluorophores were adjacent to one another. Cy3 was excited by an external 530 nm laser, while the Cy5 fluorophore was excited by the emission of the Cy3 fluorophore, allowing for observance of both fluorescent signals when the two strands were annealed to one another. Once melted, only the Cy3 emission will be observed. Binding and melting can be quantified as a function of the observance of one vs. two signals (FRET calculation, as outlined in the graphical protocol below). FRET quantifications were normalized as a value from 1 to 0 so individual experiments could be compared to one another. Cy3 and Cy5 emission intensities were assessed via a Cary Eclipse Fluorescence Spectrophotometer. Equilibrium binding was quantified by plotting normalized FRET values to protein concentration. Increasing amounts of RPA or Alt-RPA were titrated into the solution containing 1 nM of each DNA substrate, and the midpoint of DNA substrate binding was used to infer relative DNA binding affinities of RPA and Alt-RPA Protein-mediated unwinding of DNA (melting) was observed by saturating the DNA with the purified protein of interest, and observing the FRET signal over time. The time needed to reach 0.5 FRET (i.e. half of the DNA being unwound) was used to quantify the rate of substrate melting by each complex.

### FAN1 protein purification

Recombinant human FAN1 protein is expressed and purified from Sf9 insect cells as described previously (Maity et al., 2013, Deshmukh et al., 2021).

### FAN1 nuclease assay

FAN1 nuclease assays were performed in nuclease assay buffer (50 mM Tris HCl pH 8.0, 25 mM NaCl, 1 mM MnCl2, 1 mM dithiothreitol,200 mg/ml BSA) with 100 nM of fluorescently labeled DNA incubated with 50nM of FAN1 protein. Reactions were initiated by the addition of protein, incubated at 37C, for 20 minutes then stopped with formamide loading buffer (95% formamide, 10 mM EDTA). Products of were separated using 6% denaturing sequencing gel for 1 hr at 2000 V and detected at fluorescence filter in the Typhoon FLA (GE Healthcare).

### RPA and Alt-RPA subunit BioID and data analysis

#### Cloning and expression in HEK293FT cells

RPA1, RPA2, RPA3 and RPA4 expression constructs were generated from specific PCR amplification from a cDNA library, and inserted into either a pgLAP1-3MYC-BioID2 vector (RPA1, RPA2, RPA3) or a pgLAP1-FLAG-BioID2 vector (RPA4) using Gateway cloning as per the manufacture’s protocols (Gateway BP Clonase - ThermoFisher Scientific, catalogue #11789100 and Gateway LR Clonase - ThermoFisher Scientific, catalogue #11791020). This vector produces a myc-tagged construct conjugated to a functional BioID2 at the protein N-terminal. pgLAP1-3MYC-BioID2 subnit plasmids were then transformed and stably integrated into HEK293-Flp-In-T-REx cells using the manufacturer’s protocol (ThermoFisher Scientific, catalogue #R78007).

#### Biotinylation, pull down, and sample preparation

293-FT stable cell lines were induced to express pgLAP1-3MYC-BioID2-RPA 1-4 by Doxycyclin 48 hours prior to pull-down at ~40-50% cell confluency. The next day, 50 μM Biotin was added 24 hours prior to the pulldown, with or without hydroxyurea treatmen of 1 mM. After 24 hours, cells were washed 3x with cold sterile 1x PBS, trypsinised, pelleted at 1500xg for 5 minutes at 4 °C, and then washed again 2x with cold sterile 1x PBS. 600 μL of cold, freshly made lysis buffer (8M urea, 50 mM HEPES, pH 7.4, 1 mM PMSF, 1 mM DTT, 1% Triton X-100) was added to each pellet and allowed to lyse on ice for 1 hour. Pellets were then sonicated on ice (30 amplitude, 2 cycles of 10 second on and 10 seconds off, with 600 μL of fresh lysis buffer being added to the pellet between the two cycles). The lysate was centrifuged at 16,500xg for 10 minutes at 4 °C, and the supernatant was transferred to a fresh tube. Streptavidin Sepharose High Performance beads (Sigma-Aldrich, catalogue # GE17-5113-01) were washed 3x with 1 mL cold lysis buffer, and then added to each of the samples. Samples were rotated overnight with the beads at 4°C. The next day, the samples were centrifuged for 5 minutes at 1000xg, and the supernatant removed. Beads were washed 4x by rotated for 10 minutes at room temperature in 1 ml wash buffer (8M Urea, 50 mM HEPES, pH 7.4). Beads were pelleted by centrifuging for 2 minutes at 1000xg and transferred to a new tube. From this point on MS-grade water was used for the preparation of all buffers. Beads were washed 4x with 1 mL 20 mM ammonium bicarbonate water, and then 1x with the same buffer with 1 mM added biotin (to saturate unbound streptavidin). The bound proteins were then reduced using 50 μl of 20 mM ammonium bicarbonate buffer with 10 mM added DTT for 30 minutes, rotating at 60°C. Samples were cooled to room temperature for 5 minutes. Proteins were then alkylated in a light-tight container using 50 μL of 50 mM ammonium bicarbonate buffer with 15 mM chloroacetamide for 1 hour rotating at room temperature. The chloroacetamide was neutralized by adding DTT to a final concentration of 15 mM and then rotating for 3 minutes at room temperature. The proteins were then digested by adding 1 μg Pierce MS-grade trypsin (ThermoFisher Scientific, catalogue # 90058) and incubating overnight at 37°C while rotating. The next day, formic acid was added to a final concentration of 1%, and tubes rotated for 5 minutes at room temperature, to stop the reaction. Beads were centrifuged at 2000xg for 3 minutes at room temperature and the supernatant was collected into a fresh tube and then put to the side. Beads were then resuspended in 100 μL of 60% acetonitrile and 0.1% FA, and then rotated for 5 minutes at room temperature. The beads were then centrifuged again at 2000xg for 3 minutes at room temperature and the supernatant collected and added to the supernatant from two steps prior. The supernatant was dried by a centrifugal evaporator at 60°C until completely dried and then resuspended in 30 μL of 0.1% trifluoroacetic acid (TFA) buffer. Peptides were then purified with ZipTip 10-μl micropipette tips containing a C18 column as per the manufacturer’s protocol (EMD Millipore, catalogue # ZTC18M008). Peptides were eluted in new tubes, in a final volume of 30 μL comprised of 50% ACN and 1% FA buffer. The supernatant was dried by a centrifugal evaporator at 60°C until completely dried and then resuspended in 30 μL of 1% FA buffer. Peptides were then transferred to a glass vial and stored at −20°C until mass spectrometry analysis.

#### LC-MS/MS analysis

250 ng of each sample was injected into an HPLC nanoElute system (Bruker Daltonics), loaded onto a trap column with a constant flow of 4 μl/min (Acclaim PepMap100 C18 column, 0.3 mm id × 5 mm, Dionex Corporation, catalogue # 164567), and then eluted onto an analytical C18 Column (1.9 μm beads size, 75 μm × 25 cm, PepSep). Peptides were eluted over 2 hours in a gradient of acetonitrile (5-37%) in 0.1% FA at 500 nL/min while being injected into a TimsTOF Pro ion mobility mass spectrometer equipped with a Captive Spray nano electrospray source (Bruker Daltonics). Data was acquired using data-dependent auto-MS/MS with a 100-1700 m/z mass range, with PASEF enabled, number of PASEF scans set at 10 (1.27 seconds duty cycle), a dynamic exclusion of 0.4-minute, m/z dependent isolation window and collision energy of 42.0 eV. The target intensity was set to 20,000, with an intensity threshold of 2,500.

#### Protein identification by MaxQuant analysis

Raw data files were analyzed using MaxQuant version 1.6.17.0 software392 and a Uniprot human proteome database (21/03/2020, 75,776 entries). The settings used for the MaxQuant analysis (with TIMS-DDA type in group-specific parameters) were: 2 miscleavages were allowed; fixed modification was carbamidomethylation on cysteine; enzymes were Trypsin (K/R not before P); variable modifications included in the analysis were methionine oxidation, protein N-terminal acetylation and protein carbamylation (K, N-terminal). A mass tolerance of 10 ppm was used for precursor ions and a tolerance of 20 ppm was used for fragment ions. Identification values "PSM FDR", "Protein FDR" and "Site decoy fraction" were set to 0.05. Minimum peptide count was set to 1. Label-Free-Quantification (LFQ) was also selected with a LFQ minimal ratio count of 2. Both the "Second peptides" and "Match between runs" options were also allowed.

#### Data analysis and statistics

Following analysis, results were sorted by parameters set by Prostar software (Proteomics statistical analysis with R). Proteins positive for at least either one of the "Reverse", "Only.identified.by.site" or "Potential.contaminant" categories were eliminated, as well as proteins identified from a single peptide. An SLSA (Structured Least Square Adaptative) and DetQuantile imputation were performed for, respectivly, POV (Partially Observed Value) and MEC (Missing in the Entire Condition) missing values. After a mean centering within each condition, results were sorted to retain proteins that were present in at least 2 of 3 biological replicates for each condition. For the mass spectrometry analysis, the specific protein-protein interaction networks involving each of the RPA or Alt-RPA proteins was quantified based on intensities to obtained enrichment ratios and MS/MS counts. Quantification of the identified proteins for each subunit measured the enrichment in comparison to the negative control HEK293-FT. Experiments were performed in biological triplicates. Enrichment ratios where significant when over the 90% percentile of associated proteins. Enrichment ratios detected for each quantified protein were compared between subunits.

### AAV-overexpression in mouse brains

AAV vector plasmids contained cDNA for either EGFP-Rpa1 (cloned by reverse transcription and PCR of RNA from mouse primary cortical neurons) or EGFP under control of a CMV promoter. AAV vectors generated after transient transfection of HEK293T cells, and the recombinant virus was isolated from two sequential continuous CsCl gradients. AAV were injected into 5-week-old mice (randomized method for injection of a particular virus vector) into the subarachnoid space above the cerebellar surface. Mice were anesthetized intraperitoneally with Nembutal and mounted on a stereotaxic apparatus (Narishige). Forehead was tilted down 20°, and a 1 mm diameter old was made at −9.2 mm from bregma, ± 0 mm lateral to the midline. A glass syringe was inserted into the hole (along the occipital bone, 3.5 mm from the hole) and 8 μL of AAV virus solution (~1000 particles) were injected in four orientations (60, 90, 270 and 330° clockwise rotation from the posterior to anterior line, with 2 μL injected at each orientation at a rate of 0.5 ul/min).

### Fragment length analysis (capillary gel electrophoresis)

Genomic DNA was collected from mouse brain tissues following homogenisation with a MagNA Lyser Instrument (Roche; item #03358968001) (same method used in 2.3.1) and phenol:chloroform extraction with ethanol precipitation. Amplification was performed using the Expand Long Template PCR system (Roche Diagnostic, catalogue #11681834001) with 5% DMSO added. The PCR cycles were as follows: 1) 1x 95°C for 5 minutes, 2) 35x (95°C for 30 seconds, then 64°C for 30 seconds, then 72°C for 5 minutes), 3) 72°C for 10 minutes, 4) infinite hold at 4°C. PCR products were denatured with HiDi formamide and boiling at 95°C for 5 minutes, and then processed by capillary gel electrophoresis with size markers on a 3130xl Genetic Analyzer (Applied Biosystems). Peak Scanner 2 software was used to visualise the repeat sizes and repeat lengths were calculated by subtracting the length of non-repeat sequence in the PCR product and dividing by 3.

### Immunohistochemistry (IHC)

Whole mouse brains were fixed in 4% paraformaldehyde for 12-16h, embedded in paraffin, and 5 μm sagittal sections were obtained using a microtome. Xylenes was used to deparaffinize the sections, which were then rehydrated in serial dilutions of ethanol (100, 90, 80, 70%). Slides were then microwaved in 0.01M citrate buffer (pH 6.0) at 120°C for 15 min for antigen retrieval. Sections were blocked with 10% normal donkey serum in 1xPBS for 1 hour at room temperature. Sections were then incubated with primary antibodies in blocking solution for 1 hour a room temperature, washed 2x with 1xPBST, and then incubated with secondary antibodies in blocking solution for 1 hour at room temperature, washed 2x with 1xPBST, Nuclei were stained with 0.2 μg/mL DAPI in PBS (DOJINDO Laboratories, catalogue #D523), and then mounted. Images were acquired using a FV1200IX83 Olympus confocal microscope and BZ-X800 Keyence All-in-one fluorescence microscope.

## Acknowledgements

We gratefully acknowledge the National Ataxia Foundation and the University of Florida Center for NeuroGenetics for supporting and providing the donated SCA1 tissues used in the study, and the patients and family members who made this work possible. FMB is supported by the Canadian Institutes of Health Research (BMB-398925). EIC is supported by the Canadian Institutes of Health Research (PJT-159683), the Natural Sciences and Engineering Research Council of Canada (RGPIN-2016-05559), and the Garron Family Cancer Centre. ARLS is supported by the National Insitutes of Health (R35 NS122140). RJH is supported by the Hereditary Disease Foundation and holds a Berman/Topper CareerFellowship from the Huntington’s Disease Society America. DEL is supported by the Hereditary Disease Foundation. ALD is supported by the Hereditary Disease Foundation. J-YM is supported by the Canadian Institutes of Health Research (FRN-388879). CEP is supported by the Canadian Institutes of Health Research (FRN-148910; FRN-173282), the Natural Sciences and Engineering Research Council of Canada (RGPIN-2016-08355 RGPIN-2016-06355/498835), The Petroff Family Foundation, Tribute Communities, The Marigold Foundation, The Kazman Family Foundation. J-YM. holds a Tier 1 Canada Research Chair in DNA Repair and Cancer Therapeutics. CEP holds a Tier 1 Canada Research Chair in Disease-Associated Genome Instability.

